# Endothelial-derived sphingolipids are required for vascular development and systemic lipid homeostasis

**DOI:** 10.1101/2022.03.22.485296

**Authors:** Andrew Kuo, Antonio Checa, Colin Niaudet, Bongnam Jung, Zhongjie Fu, Craig E. Wheelock, Sasha A. Singh, Masanori Aikawa, Lois E. Smith, Richard L. Proia, Timothy Hla

## Abstract

Serine palmitoyl transferase (SPT), the rate-limiting enzyme in the *de novo* synthesis of sphingolipids (SL), is needed for embryonic development, physiological homeostasis, and response to stress. The functions of *de novo* SL synthesis in vascular endothelial cells (EC), which line the entire circulatory system, are not well understood. Here we show that the EC *de novo* synthesis not only impacts vascular development but also maintains normal SL metabolic homeostasis in the circulatory system and peripheral organs. Mice with an endothelial-specific gene knockout of SPTLC1 (*Sptlc1* ECKO), an essential subunit of the SPT complex, exhibited EC-intrinsic effects including reduced EC proliferation and tip/stalk cell differentiation, resulting in delayed retinal vascular development. In addition, *Sptlc1* ECKO mice had reduced pathological retinal neovascularization in the oxygen-induced retinopathy model, suggesting that EC SL produced from the *de novo* pathway are needed for efficient VEGF signaling within the vascular system. Post-natal deletion of the EC *Sptlc1*also showed cell-extrinsic effects, including rapid reduction of several SL metabolites in plasma, red blood cells and peripheral organs (lung and liver) but not in the retina, part of the central nervous system (CNS). In the liver, EC *de novo* SL synthesis was required for acetaminophen-induced ceramide elevation and hepatotoxicity. These results suggest that EC-derived SL metabolites are in constant flux between the vasculature, circulatory elements, and parenchymal cells of non-CNS organs. Taken together, our data point to the central role of the endothelial SL biosynthesis in maintaining vascular development and neovascular proliferation, non-CNS tissue metabolic homeostasis and hepatocyte response to stress.

## Introduction

Sphingolipids (SL), which comprise ∼10-20 % of total cellular lipids, are involved in wide range of biological processes. Complex membrane SL, sphingomyelin (SM) and glycosphingolipids, are enriched in lipid rafts and caveolae, which provide an optimal platform for intracellular signaling and cell-cell interactions (van Meer *et al*, 2008). In contrast, the secreted SL, sphingosine-1-phosphate (S1P), acts as a ligand for G-protein coupled receptors (S1PR1-5) which regulate many physiological and pathological functions (Proia & Hla, 2015). While SL can be derived from dietary sources, most cellular SL are synthesized via the *de novo* pathway. This pathway is initiated in the endoplasmic reticulum by the condensation of serine and fatty acyl-CoA to generate 3-ketosphinganine by the serine palmitoyltransferase (SPT) enzyme complex. The product, 3-ketosphinganine, is further converted into dihydrosphingosine, dihydroceramide, and ultimately ceramide (Cer), which is a substrate for the production of complex SL and metabolites such as sphingosine (Sph) and S1P (Merrill, 2002). The SPT complex, which is composed of long chain subunit 1 (SPTLC1), subunit 2/3 (SPTLC2/3) and small subunit A/B (SPTSSA/B) subunits, catalyzes the rate-limiting step in the *de novo* SL biosynthetic pathway (Han *et al*, 2009). Absence of any of the subunits abolishes the enzymatic activity of SPT, which is required for normal embryonic development in mice (Hojjati *et al*, 2005). In addition, SPT enzyme activity is controlled by interaction with ORMDL proteins, which respond to extracellular and cell-intrinsic cues to regulate the flux of SL synthesis by the *de novo* pathway (Siow *et al*, 2015) (Han *et al*, 2019).

The significance of *de novo* SL biosynthetic pathway in organ function and diseases have been deduced from genetic association studies and mouse genetic models. Mutations in the *Sptlc1* gene are associated with peripheral neuronal dysfunction including hereditary sensory neuropathy type 1 (HSAN1) (Bejaoui *et al*, 2001) (Dawkins *et al*, 2001), childhood amyotrophic lateral sclerosis (ALS) (Mohassel *et al*, 2021) and macular telengectasia type-2 (Gantner *et al*, 2019). *Sptlc2* haploinsufficiency in the macrophages causes reduction of circulating SM and enhancement of reverse cholesterol transport in murine genetic models (Chakraborty *et al*, 2013), suggesting the requirement of this pathway in maintaining systemic SL homeostasis. Loss of function of the SPT complex in the hepatocyte reduces SM in the plasma membrane and greatly reduces cadherin, an important protein for adherens junctions (Li *et al*, 2016), highlighting the importance of this pathway in maintaining proper plasma membrane function. Collectively, these data demonstrate cell type-specific function for *de novo* SL synthesis.

Endothelial cells (EC) line the inner layer of the entire circulatory system and mediate vascular tone control, barrier integrity, oxygen supply, immune cell trafficking and waste removal. The vasculature of the central nervous system (CNS), which includes the brain and retina, is unique in that EC help form a highly selective barrier for circulatory metabolites. Indeed, EC in CNS forms a specialized structure between the blood and the tissue, known as the blood brain barrier (BBB) and blood retinal barrier (BRB) that consist of cellular compartments including EC, pericytes, and neural cells (astrocytes, glia and neurons). SL metabolites, originally isolated from brain due to their relative abundance, are critical for CNS development, physiology and disease (Olsen & Faergeman, 2017). Metabolism of lipids are compartmentalized in CNS and non-CNS tissues, as it has been shown for cholesterol (Orth & Bellosta, 2012). Whether SL metabolism in CNS and non-CNS organs is compartmentalized and the role of SL in organotypic vascular beds is not known.

Our current knowledge about SL metabolism in EC has been centered around S1P, a key lipid mediator needed for developmental and pathological angiogenesis, and Cer, a mediator of stress responses (Cartier & Hla, 2019) (Jernigan *et al*, 2015). Activation of S1P signaling in the EC S1PR1 suppresses vascular endothelial growth factor (VEGF) dependent vascular sprouting and maintains barrier integrity, resulting in stable and mature vessels (Jung *et al*, 2012) (Gaengel *et al*, 2012). Upon irradiation, Cer aggregate to form Cer-enriched membrane domains within the cell membrane, which regulate physicochemical properties of membranes, leading to apoptosis and capillary leak (Kolesnick & Fuks, 2003). Moreover, VEGFR2 has been shown to colocalize with SL enriched lipid rafts on the plasma membrane and impact VEGF induced intracellular signal transduction processes (Laurenzana *et al*, 2015). These results suggest that multiple SL metabolites regulate vascular development (angiogenesis), homeostasis and diseases. Recently, it has been shown that EC-specific deletion of *Sptlc2* results in lower blood pressure and reduced plasma Cer, implying the protective function of *de novo* SL biosynthetic pathway in vascular disease (Cantalupo *et al*, 2020). However, the role of *de novo* SL biosynthetic pathway in metabolic flux of SL metabolites in various tissue compartments and vascular development is not known.

To address the questions above, we generated a mouse model in which *de novo* synthesis of SL is abolished by *Sptlc1* gene deletion in endothelial cells (*Sptlc1* ECKO). *Sptlc1* ECKO mice exhibited delayed retinal vascular development and reduced pathological angiogenesis in an oxygen-induced retinopathy model. Moreover, *Sptlc1* ECKO mice showed rapid decline of many SL, including dihydrosphingosine (dhSph), dihydrosphingosine-1-phosphate (dhS1P) and Cer in plasma and red blood cells (RBC), demonstrating SL flux from EC to circulation. SL content of peripheral tissues such as lung and liver were also reduced upon *Sptlc1* EC deletion. Reduced Cer levels in *Sptlc1* ECKO mice protected against acetaminophen-induced hepatoxicity. Collectively, our data not only demonstrate the role of *de novo* synthesis of SL during vascular development, but also reveal a novel function of EC as a source to provide SL to the circulation and to peripheral tissues for their proper functions and response to stress.

## Results

### Endothelial SL are reduced after tissue-specific deletion of *Sptlc1* gene

We generated a mouse strain in which *Sptlc1* floxed mice are intercrossed with tamoxifen-inducible VE-cadherin Cre recombinase mice (*Sptlc1* ECKO, Figure 1-figure supplement 1A). Mice were administered tamoxifen at 6-week of age and followed for 10 weeks. We did not observe any difference in weight gain or any gross abnormalities (Figure 1-figure supplement 1B). Blood chemistry analyses indicated that *Sptlc1* ECKO mice have normal liver enzyme levels, kidney function and blood chemistry (Figure 1-figure supplement 1C). To determine the efficiency and specificity of *Sptlc1* gene deletion, we prepared EC (CD31^+^; CD45^-^) and non-EC (CD31^-^) from single-cell suspensions isolated from enzymatically digested lung tissues (Figure 1A). Flow cytometry analyses showed that isolated EC population was ∼90% pure and non-EC population contains < 10% of contaminating CD31^+^ cells (Figure 1B). SPTLC1 expression was reduced markedly (∼80%) in *Sptlc1* ECKO endothelium specifically (Figure 1C-E). In contrast, SPTLC1 expression in tissues with lower EC abundance such as bone marrow did not show differences (Figure 1-figure supplement 2). These results demonstrate that SPTLC1 abundance is abolished in a cell-specific manner in *Sptlc1* ECKO mice.

**Figure 1.**
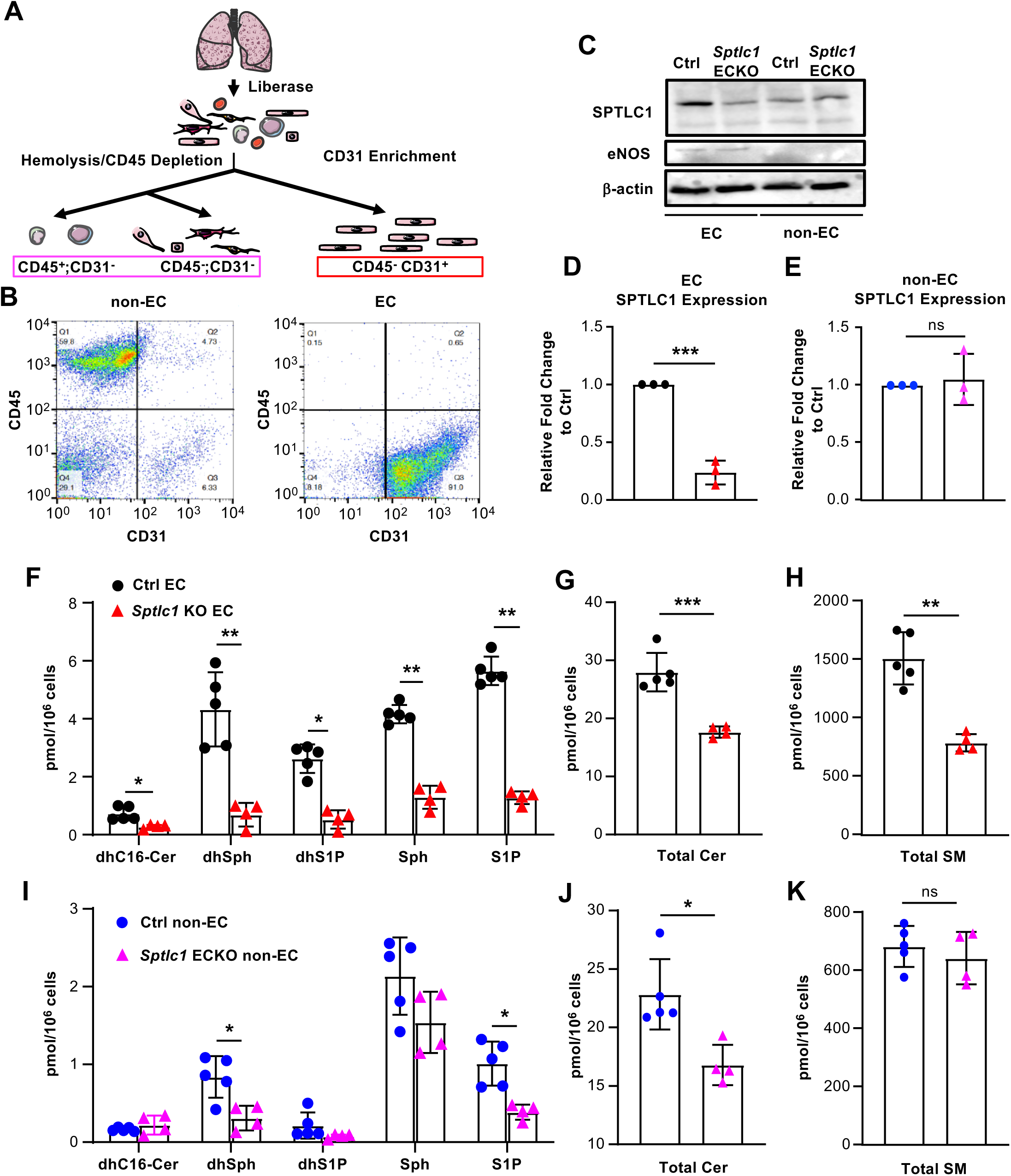
Ablation of *Sptlc1* in EC reduces sphingolipid content in both EC and non-EC cell population of the lung. (A) Scheme of lung EC and non-EC isolation procedure used in this figure. (B) Flow cytometry analysis for determination of lung Non-EC and EC population purities. Non-EC is identified as CD45+/-; CD31- cells. EC is identified as CD45-; CD31+ cells. (C) EC and non-EC from Ctrl and *Sptlc1* ECKO lungs were analyzed by Western Blotting analysis for SPTLC1 and eNOS expressions. β-actin was used as an internal control. Relative fold change of SPTLC1 to control in EC and non-EC population were quantified as shown in (D) an (E). SL content in Ctrl and *Sptlc1* KO EC were determined by LC-MS/MS, including dhC16-Cer, dhSph, dhS1P, Sph and S1P in (F), total Cer content in (G), and sphingomyelin (SM) in (H). SL content in Ctrl and *Sptlc1* KO non-EC were determined by LC-MS/MS, including dhC16-Cer, dhSph, dhS1P, Sph and S1P in (I), total Cer content in (J), and total SM content in (K). Data are expressed as mean±SD. Statistical significance was determined by unpaired t test. *p < 0.05; **p < 0.01. ***p<0.001. ns, non-significant.

We next measured SL metabolites in isolated lung EC from *Sptlc1* ECKO mice by LC-MS/MS. SL species in the *de novo* biosynthetic pathway, namely, dhSph, C16-dihydroceramide (dhC16-Cer) were markedly reduced (Figure 1F). In addition, Sph, S1P and dhS1P were reduced. Cer species, including C16:0, C22:1, C22:0, C24:0, C24:1 and C26:0 were also reduced, resulting in decreased total EC Cer levels (Figure 1G, Figure 1-figure supplement 3A). Moreover, SM species, including C16:0, C22:0, C24:0, C26:1, C26:0 SM (Figure 1H, Figure 1-figure supplement 3B) were reduced, resulting in decreased total EC SM content. These data show that lack of SPTLC1 subunit leads to marked reduction in EC SL metabolites.

We noticed that dhSph, S1P and several Cer species were significantly reduced in non-EC populations of *Sptlc1* ECKO while some SL (dhC16-Cer, dhS1P, Sph and SM) were unaffected (Figure 1I-K, Figure 1-figure supplement 3C and D). Given that non-EC population contained less than 10% of CD31+ cells (Figure 1B), the reduction is likely not due to EC contamination. This surprising finding suggests that EC-derived SL metabolites are transferred to non-EC parenchymal cells in the lung.

### EC SPT regulates retinal vascular development and disease

We next investigated whether EC SPT is important for vascular development. Vascular development occurs in the murine retina after birth and is fully formed by postnatal day 30 (P30) (Rust *et al*, 2019). *Sptlc1* ECKO mice exhibited a lower retinal vascular density and delayed radial expansion of retinal vascular network at P6 compared to controls (Figure 2A-C) while pericyte coverage was not altered (Figure 2D and E). Proliferation of retinal EC was reduced in *Sptlc1* ECKO mice as indicated by phosphorylated histone H3 staining (Figure 2F and G). Tip cell number, which require the function of VEGF on developing vascular network, was markedly reduced in *Sptlc1* ECKO mice, as indicated by ESM1 staining of developing retina (Figure 2F and H). Reduced tip cell formation and proliferation of stalk cells likely resulted in the abnormal retinal vascular development. The deeper vascular plexus, which begins to form ∼ P9 by vertical expansion of vascular sprouts from the superficial vascular network, also showed marked developmental delay in *Sptlc1* ECKO mice compared to controls (Figure 2I-K). Expressions of EC proteins important for BRB formation, including LEF1 (a key transcription factor involved in canonical Norrin/Wnt signaling), TFRC (transferrin receptor C), and CLDN5 (Claudin-5, a tight junction protein) were not altered at P6, suggesting that retinal EC organotypic specialization is not affected in the absence of SPT (Figure 3A-E). Interestingly, abundance of MFSD2A, a lysophospholipid transporter that suppresses transcytosis in EC, were upregulated in the retinal arteries of *Sptlc*1 ECKO retina (Figure 3D and F). The delayed vascular developmental phenotype was normalized at P15 (Figure 3-figure supplement 1). We surmise that SL synthesized by other cells of the retina or derived from circulatory sources were used to rescue the *Sptlc1* ECKO phenotype in the vasculature. Together, these results suggest that active SPT function in the endothelium is needed for normal vascular development.

**Figure 2.**
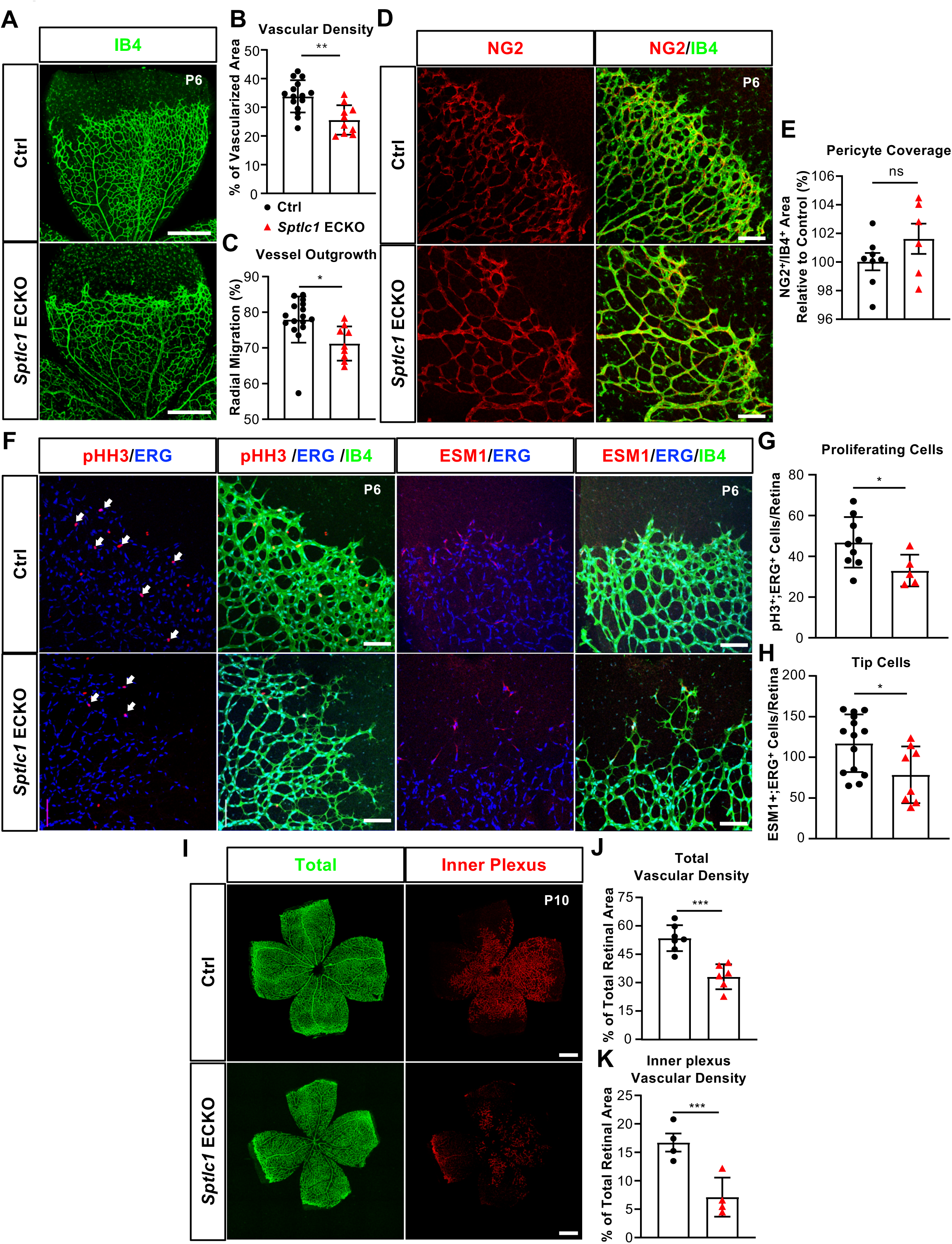
Retinal vascular development is delayed in the absence of *Sptlc1* in EC. (A) Retinal vascular plexus at P6 of Ctrl and *Sptlc1* ECKO mice were immunostained with Isolectin-B4 (IB4). Vascular density and outgrowth were quantified in (B) and (C). (D) Pericytes were immunostained with NG2 and pericytes coverage was quantified by NG2^+^/IB4^+^ area as shown in (E). (F) Proliferating cells and tip cells mice were immunostained with phospho-histone H3 (pHH3) and ESM1 respectively. ERG immunostaining is served as an EC marker. White arrows indicate pHH3^+^/ERG positive cells and quantified in (G). Tip cells were quantified by ESM1^+^/ERG^+^ cells as shown in (H). (I) Retinal vascular plexus at P10 of Ctrl and *Sptlc1* ECKO mice were immunostained with IB4. Representative images of total and inner plexus were shown and quantified in (J) and (K). (L) Total vasculature at P15 of control and *Sptlc1* ECKO mice were immunostained with IB4 and quantified in (M). Data are expressed as mean±SD. Statistical significance was determined by unpaired t test. *p < 0.05; **p < 0.01; ***p < 0.001. ns, nonsignificant. Scale bar in (A): 500μm, in (D) and (F)): 100μm, in (I) and (L): 500μm

**Figure 3.**
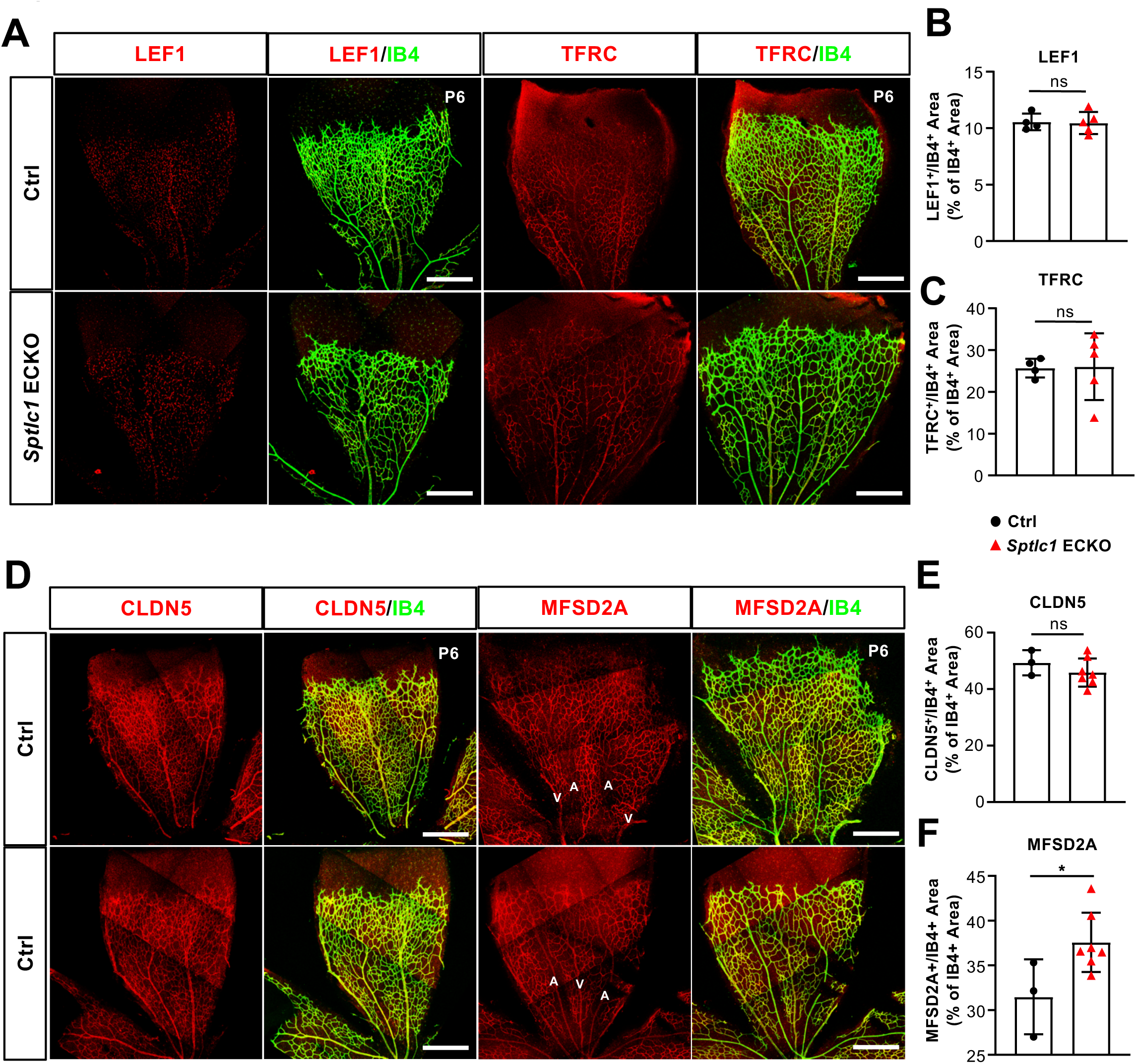
Blood-retina barrier (BRB) protein expressions are not reduced in the retina of *Sptlc1* ECKO mice. (A) BRB specific proteins including LEF1 and TFRC in Ctrl and *Sptlc1* ECKO P6 retina were immunostained and quantified by colocalization with IB4 staining shown in (B) and (C). (D) Tight junction protein, CLDN5 in Ctrl and *Sptlc1* ECKO P6 retina were immunostained and quantified by colocalization with IB4 staining shown in (E). BRB specific protein, MFSD2A, in Ctrl and *Sptlc1* ECKO P6 retina were immunostained and quantified by colocalization with IB4 staining shown in (F). Arterial and venous regions are marked as “A” and “V”. Data are expressed as mean±SD. Statistical significance was determined by unpaired t test. *p < 0.05; ns, non-significant. Scar bars, 100μm.

Oxygen-induced retinopathy (OIR) in mice is a widely used disease model to investigate pathological neovascularization during retinopathy of prematurity, diabetic retinopathy and wet form of age-related macular degeneration (Smith *et al*, 1994) (Campochiaro, 2013). VEGF produced by Müller glia and astrocytes is a major contributor to pathological vascular tuft formation in this model (Weidemann *et al*, 2010). We used this OIR model in WT and *Sptlc1* ECKO mice to determine the role of EC-derived SL metabolites in pathological angiogenesis. Tamoxifen was administered at P1- P3 to induce *Sptlc1* deletion and mouse pups were exposed to 75% O_2_ from P7-P12 (Figure 4A). This hyperoxia environment causes capillary dropout (vaso-obliteration) due in part to suppression of VEGF production (Smith *et al*., 1994). In the absence of EC *de novo* SL synthesis, vascular dropout was exacerbated at P10 and P12, resulting in reduced vascular network density and increased avascular areas (Figure 4B-G). These results suggest that EC SL *de novo* synthesis impacts VEGF sensitivity of the developing vasculature. Upon return to normoxia, avascular zones are expected to be severely hypoxic. To examine pathological vascular tuft formation, we induced EC *Sptlc1* deletion at P12 which is the beginning of the retinal hypoxia in this animal model (Fig. 4H). With this timed gene deletion strategy, we observed significant attenuation of pathological neovascularization in *Sptlc1* ECKO mice at P17 (Figure 4I and J). In addition, vaso-obliterated regions were larger in *Sptlc1* ECKO mice (Figure 4I and K). Collectively, the results from the OIR model indicated that *de novo* synthesis of SL in EC determine the pathological phenotypes in the retinal vasculature (vessel dropout and pathological vascular tuft formation), likely due to reduced VEGF responsiveness.

**Figure 4.**
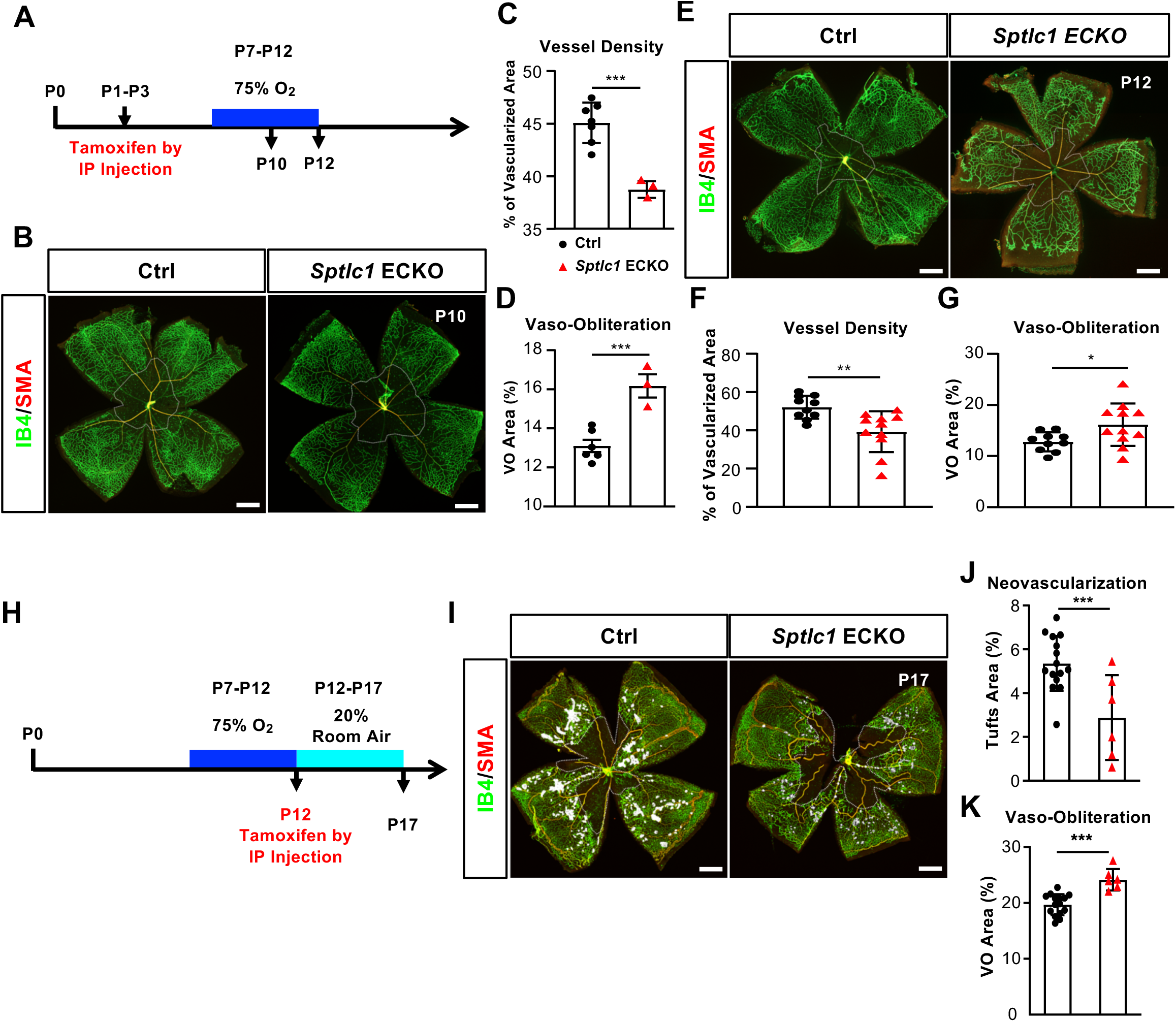
*Sptlc1* ECKO mice show enhanced vaso-obliteration and reduced neovascularization in oxygen induced retinopathy model. (A) Schematic of timeline for tamoxifen treatment, oxygen challenge and tissue collection used from (B) to (G). (B) P10 retinas of control and *Sptlc1* ECKO mice were immunostained with IB4 and smooth muscle actin (SMA). Vascular density and vaso-obliteration (dashed line area) were quantified in (C) and (D). (E) P12 retinas of control and *Sptlc1* ECKO mice were stained with IB4 and SMA. Vascular density and vaso-obliteration (dashed line area) were quantified in (F) and (G). Avascularized area is highlighted in dashed line. (H) Schematic of timeline for tamoxifen treatment, oxygen challenge and tissue collection used from (I) to (K). Note that tamoxifen was administered after 75% O2 challenge. (I) P17 retinas of control and *Sptlc1* ECKO mice were immunostained with IB4 and SMA. Neovascularization (white area) and vaso-obliteration (dashed line area) were quantified in (J) and (K) Data are expressed as mean±SD. Statistical significance was determined by unpaired t test. *p < 0.05; **p<0.01, ***p < 0.001. ns, non-significant. Scale bars, 500μm.

### Endothelial SPT activity is critical to maintain circulatory SL metabolic homeostasis

Since the non-EC parenchymal cells isolated from the lungs of *Sptlc1* ECKO mice contain reduced levels of SL metabolites (Figure 1I-K), we hypothesized that EC-derived SL are transferred to non-EC parenchymal cells either directly (cell-cell transfer) or indirectly via the circulatory system. We first quantified SL metabolites in plasma isolated from adult *Sptlc1* ECKO mice. S1P an dhS1P levels were reduced ∼60% in *Sptlc1* ECKO (Figure 5A). This reduction is more pronounced than that observed in sphingosine kinase-1 and −2 ECKO (Gazit *et al*, 2016) and S1P transporter (SPNS2) ECKO mice (Hisano *et al*, 2012; Mendoza *et al*, 2012). dhSph, a sphingoid base synthesized in the *de novo* synthesis pathway, was also reduced significantly (Figure 5A). Moreover, Cer species (C24:1, C22:0, C26:1 and C26:0) and C 24:0 SM were reduced in *Sptlc1* ECKO plasma, which resulted in reduction of total Cer in plasma (Figure 5B-E). These results indicate that SL *de novo* synthesis pathway in EC is a significant source of sphingoid bases, their phosphorylated metabolites and Cer in plasma.

**Figure 5.**
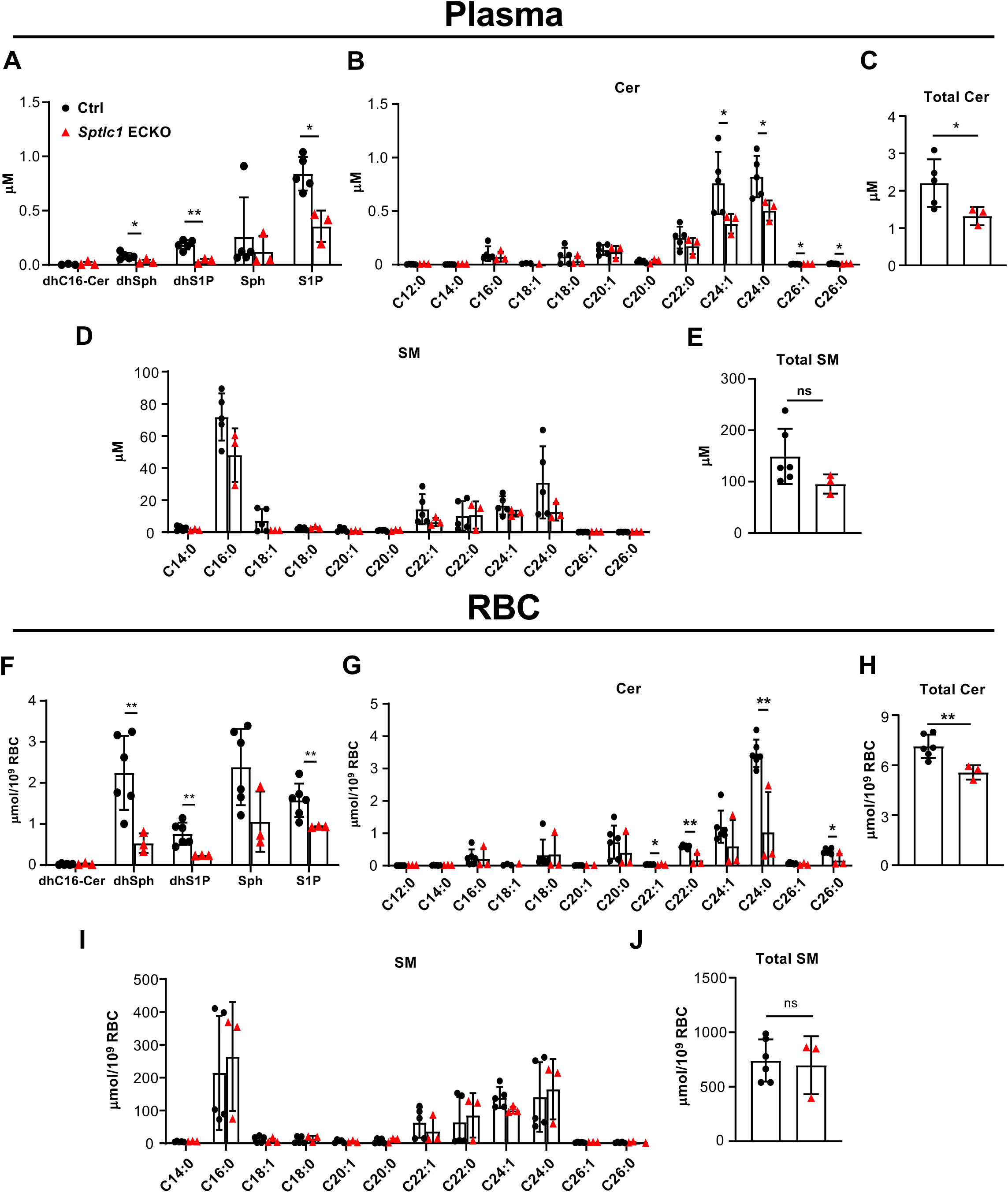
*Sptlc1* ECKO displays reduced SL content in circulation. SL content in plasma from Ctrl and *Sptlc1* ECKO mice was determined by LC-MS/MS, including dhC16-Cer, dhSph, dhS1p, Sph and S1P in (A), Cer with different fatty acyl chain length in (B), total Cer content in (C), and SM with different fatty acyl chain length in (D), and total SM in (E). Sphingolipid content in RBC from Ctrl and *Sptlc1* ECKO mice was determined by LC-MS/MS, including dhC16-Cer, dhSph, dhS1p, Sph and S1P in (F), Cer with different fatty acyl chain length in (G), total Cer content in (H), and SM with different fatty acyl chain length in (I), and total SM in (J). Data are expressed as mean±SD. Statistical significance was determined by unpaired t test. *p < 0.05; **p < 0.01. ns, non-significant.

RBC, which store and release S1P, are a major contributor of plasma S1P (Pappu *et al*, 2007). We therefore quantified SL metabolites in RBC of *Sptlc1* ECKO mice. Sphingoid bases and their phosphorylated counterparts (dhSph, dhS1P, and S1P) were significantly reduced in *Sptlc1* ECKO RBCs (Fig. 5F). Cer species (C22:1, C22:0, C24:0 and C26:0) were also reduced (Figure 5G and H). In contrast, SM species were not altered in RBC of *Sptlc1* ECKO mice (Figure 5I and J). These results show that reduction of SL metabolites in EC result in concomitant changes in plasma and RBC pools of SL metabolites. This suggests the existence of a SL metabolic flux from EC to circulatory compartments.

We next determined the kinetics of SL reduction in plasma/RBC after the deletion of EC *Sptlc1*. Tamoxifen was injected at 6 weeks of age to induce deletion of EC *Sptlc1* and plasma/RBC SL metabolites were temporally quantified. Rapid reduction of plasma S1P levels observed as early as 1 week and maintained thereafter (Figure 6A). Kinetics of RBC S1P reduction was similar (Figure 6B), suggesting that EC SL metabolites are rapidly secreted into plasma and that RBC and plasma SL pools are in homeostatic equilibrium.

**Figure 6.**
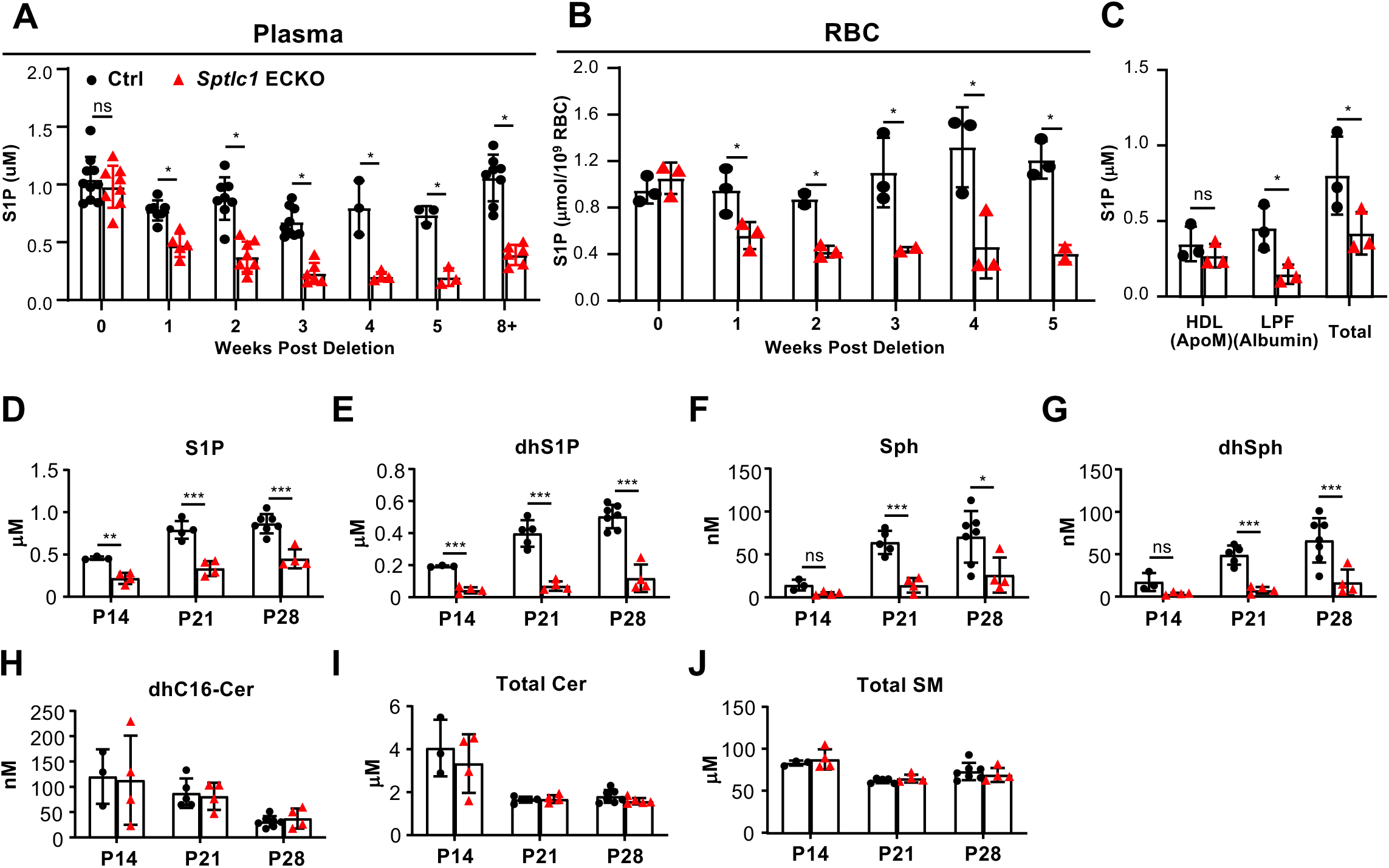
Rapid reduction of SL content in circulation upon *Sptlc1* deletion. S1P levels from plasma and RBC of Ctrl and *Sptlc1* ECKO mice were measured upon *Sptlc1* deletion over time as shown in (A) and (B). (C) Plasma from Ctrl and *Sptlc1* ECKO mice were fractionated by HPLC into HDL (ApoM enriched), Lipoprotein Free (LPF) (Albumin enriched). S1P levels in each fraction, including total plasma were measured by LC-MS/MS. Sphingolipid content from plasma samples of Ctrl and postnatal (P1-P3) deletion of *Sptlc1* in EC were measured at P14, P21 and P28 by LC-MS/MS. Species measured are as followed: S1P (D), dhS1P (E), Sph (F), dhSph (G), dhCer (H), total Cer (I), and total SM (J). Data are expressed as mean±SD. Statistical significances in (A) and (B) were determined by Two-way ANOVA test. Statistical significances in (C)-(G) were determined by unpaired t test. *p < 0.05; **p < 0.01; ***p<0.001. ns, non-significant.

Plasma S1P is bound to two chaperones, namely, Apolipoprotein M (ApoM) on HDL and albumin in the lipoprotein free fraction (LPF) (Murata *et al*, 2000). ApoM and albumin levels were not altered in *Sptlc1* ECKO plasma (Figure 6-figure supplement 1). By fractionation of plasma using fast protein liquid chromatography, we determined that ApoM bound S1P was not affected while albumin bound S1P in the LPF was significantly reduced in *Sptlc1* ECKO mice (Figure 6C). Since albumin bound S1P is less stable (Fleming *et al*, 2016), released S1P from RBC or EC first bind to albumin, followed by stable interaction with ApoM. Nevertheless, these data suggest that EC SPT-derived S1P is secreted into plasma and equilibrates rapidly with RBC storage pools.

To determine whether the *de novo* synthesis of SL in EC also contributes to circulatory pools of SL species in early postnatal mice, we deleted EC *Sptlc1* from P1-P3 followed by measurement of plasma SL metabolites. Sphingoid bases and phosphorylated derivatives (dhSph, Sph, S1P, dhS1P) showed significant reduction from P14 to P28 in *Sptlc1* ECKO mice (Figure 6D-G). In contrast, plasma dhCer, Cer and SM were not changed (Figure 6H-J, Figure 6-supplement 2). These results suggest that EC actively provide SL metabolites to circulation postnatally. Differential reduction in SL metabolites in plasma could be due to contributions from non-EC sources, such as the diet and metabolic flux from other cell types at different ages.

### *De novo* synthesis of SL in EC impacts peripheral organ SL content

We examined whether EC SPT enzyme loss affects SL metabolite levels in various organs. We analyzed the liver, lung, and retina because these organs are supplied by vessels with specialized organotypic endothelium (Potente & Makinen, 2017). The liver contains sinusoidal EC while lung vasculature is lined by EC specialized for gas exchange. In contrast, the retinal vasculature is lined by CNS-type endothelium which contains numerous tight junctions and specialized transporters with minimal transcytotic vesicles (Andreone *et al*, 2017). DhSph, a direct product of the *de novo* SL biosynthetic pathway was significantly reduced in lung and liver of *Sptlc1* ECKO mice (Figure 7A and D). In contrast, S1P and Sph levels in these tissues were not reduced (Figure 7A and D). Cer content was reduced about 30-50% in both organs from *Sptlc1* ECKO mice with C24:0 and C24:1 Cer species exhibiting largest changes (Figure7 B and C, E and F). These results suggest that EC supply SL metabolites from the *de novo* pathway into circulation which then enters liver and lung SL pools. We also measured SL content of retina, a CNS organ system. In sharp contrast to lung and liver, none of the SL metabolites were altered by lack of EC SPT (Figure 7G-H). These results reveal that the SL metabolic flux from EC to adjacent tissues only occurs in circulation and non-CNS organs such as the lung and the liver.

**Figure 7.**
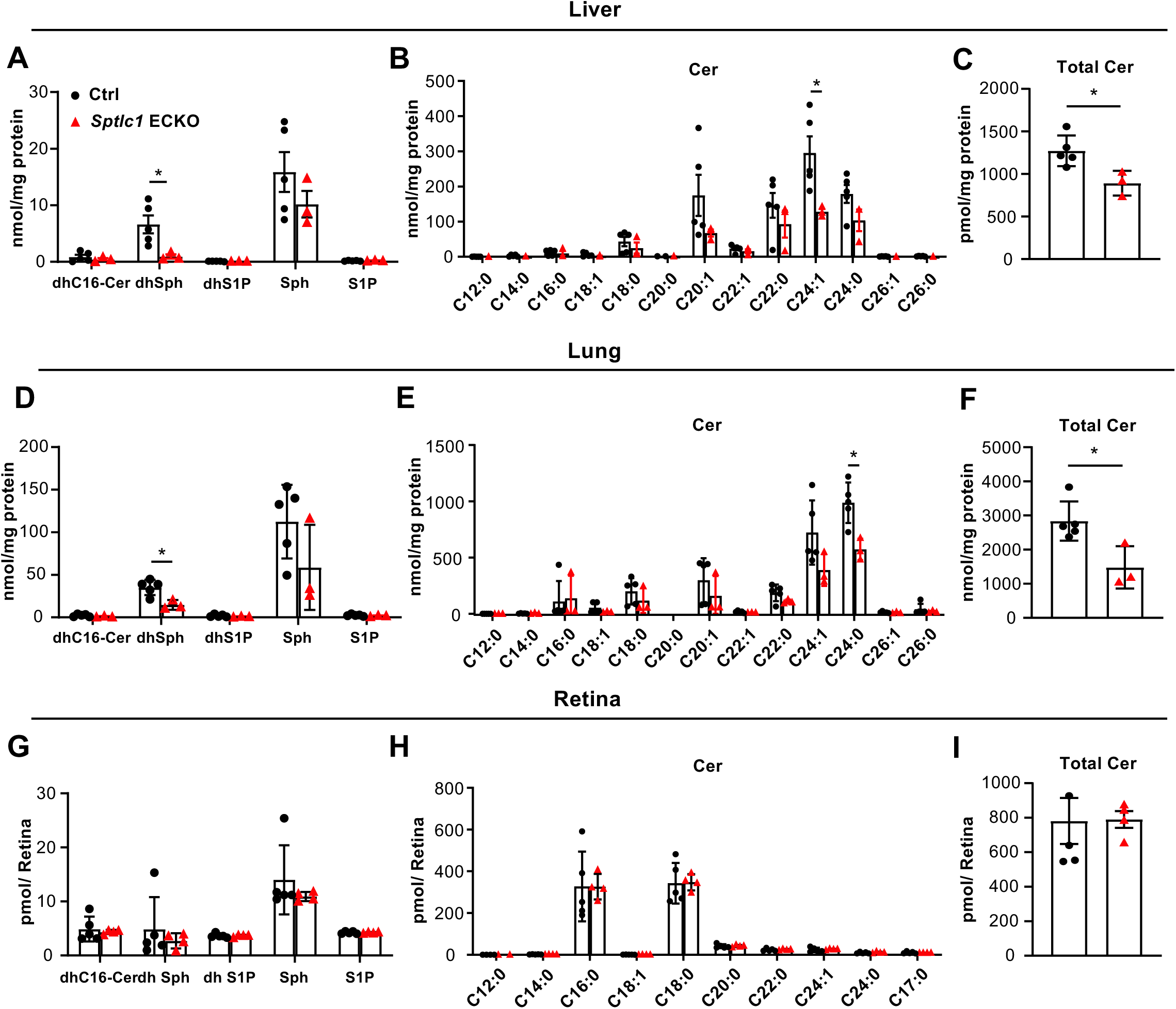
*Sptlc1* ECKO mice exhibit reduced SL content in peripheral but not CNS organs. SL content from adult liver homogenate of Ctrl and *Sptlc1* ECKO mice were measured by LC-MS/MS, including dhCer, dhSph, dhS1P, Sph and S1P (A), Cer with different fatty acyl chain length (B) and total Cer (C). SL content from adult lung homogenate of Ctrl and *Sptlc1* ECKO mice were measured by LC-MS/MS, including dhCer, dhSph, dhS1P, Sph and S1P (D), Cer with different fatty acyl chain length (E) and total Cer (F). SL content from P28 retina homogenate of Ctrl and *Sptlc1* ECKO mice were measured by LC-MS/MS, including dhCer, dhSph, dhS1P, Sph and S1P (G), Cer with different fatty acyl chain length (H) and total Cer (I). Data are expressed as mean±SD. Statistical significance was determined by unpaired t test. *p < 0.05.

### *Sptlc1 ECKO* mice are resistant to acetaminophen-induced liver injury

Liver function was normal in *Sptlc1* ECKO suggesting that reduced SL metabolites are not critical for hepatic homeostasis. However, chemical stress elevates SL metabolites such as Cer, which are involved in acute liver injury (Li et al., 2020). Alternatively, changing the sphingolipid acyl chain length composition compromises gap junction in hepatocyte, serving as a mechanism for SL regulating drug-induced liver injury (Park *et al*, 2013). To determine if EC derived SL influence hepatocyte function, we examined hepatotoxicity induced by N-acetyl-para-aminophenol (APAP; a.k.a. Acetaminophen/Tylenol) in *Sptlc1* ECKO mice. APAP treatment resulted in reduced liver injury (aspartate aminotransferase (AST) levels and liver necrosis) in *Sptlc1* ECKO mice compared with controls at 4 and 8 h following drug administration. Equivalent degrees of injury were seen in both cohorts at 24 h post-APAP (Figure 8A-C). Liver SL metabolites were quantified at 8 and 24 h after APAP administration. Many SL metabolites (dhCer C16:0, dhSph, dhS1P, Sph, S1P, Cer C24:1 and total Cer) were lower in the liver at 8 h post-APAP but only dhCer C16:0 and dhSph were lower in the *Sptlc1* ECKO mice at 24 h (Figure 8D-I). This is consistent with Cer species induced by APAP as hepatotoxicity ensued. These data implicated that reduced Cer levels in the liver of *Sptlc1* ECKO mice at 8 h may be responsible for resistance to APAP-induced liver injury at early time points.

**Figure 8.**
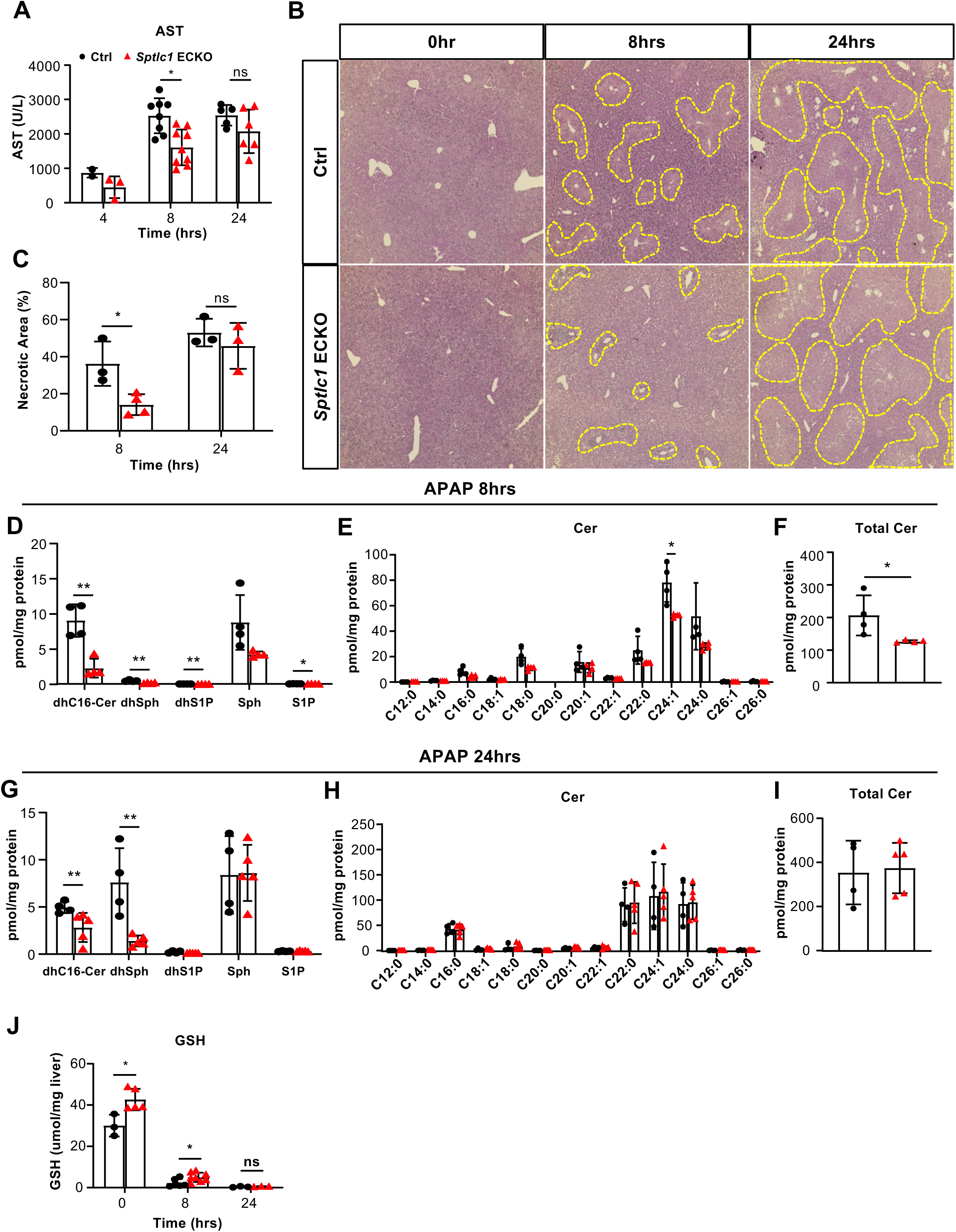
*Sptlc1* ECKO mice are protected against APAP-induced hepatoxicity. AST levels were measured from plasma samples of Ctrl and *Sptlc1* ECKO mice after 4, 8 and 24 hours of APAP injection. (B) Hematoxylin and eosin staining of Ctrl and *Sptlc1* ECKO liver after 0, 8 and 24 hours of APAP administration. Dotted lined areas indicate necrotic region and the percentage of necrotic areas were quantified in (C) (each data point represents average of 5 images from one mouse). SL content, including dhC16-Cer, dhSph, dhS1P, Sph and S1P (E), Cer with different fatty acyl chain length (F), total Cer content (G) were measured in Ctrl and *Sptlc1* ECKO liver after 8hrs of APAP injection was measured by LC-MS/MS. SL content, including dhC16-Cer, dhSph, dhS1P, Sph and S1P (H), Cer with different fatty acyl chain length (I), total Cer content (J) were measured in Ctrl and *Sptlc1* ECKO liver after 24 hours of APAP injection was measured by LC-MS/MS. (D) GSH levels were measured from liver samples of Ctrl and *Sptlc1* ECKO mice after 0, 8 and 24 hours of APAP injection. Data are expressed as mean±SD. Statistical significance was determined by unpaired t test. *p < 0.05. **p<0.01. Scale bar, 1mm

To determine the mechanisms involved in early protective response to APAP toxicity in *Sptlc1* ECKO mice, we examined levels of glutathione (GSH), the main antioxidant that protects from APAP-induced hepatoxicity. Liver GSH levels were elevated *Sptlc1* ECKO mice prior to APAP injection. Even though GSH is diminished after APAP-administration, Sptlc*1* ECKO mice still had higher GSH levels than the WT counterparts at 8hrs (Figure 8J). Thus, reduced flux of SL metabolites from EC to liver results in attenuated drug-induced hepatotoxicity via modulation of antioxidant GSH levels.

## Discussion

Lipids are derived both from the diet and from biosynthetic routes in organs. Their concentrations in various cells and tissues are regulated stringently to maintain normal physiological functions. The numerous SL species are no exception. Altered SL levels due to mutations in biosynthetic or degradative enzymes lead to embryonic lethality or severe genetic diseases (Hannun & Obeid, 2018). However, SL homeostasis in physiology of various organ systems is poorly understood. Differentiated cell types which supply SL for systemic metabolic needs and intrinsic signal transduction events are not well studied. In this study, we show that *de novo* SL biosynthesis in EC regulates developmental and pathological angiogenesis. We also discover a novel function of EC as a major SL source into circulation, which impacts SL content of peripheral tissues and the ability to withstand stress in these organs.

The SPT enzyme complex catalyzes the first committed step in the *de novo* SL biosynthetic pathway. This step is highly regulated by ORMDL proteins and other poorly understood mechanisms (Cantalupo *et al*, 2015; Linn *et al*, 2001; Siow *et al*., 2015). Since the vascular EC traverse all organ systems, we developed and analyzed a mouse genetic model in which S*ptlc1*, encoding the non-redundant subunit of SPT complex (SPTLC1) is disrupted in a tissue-specific manner using tamoxifen-induced gene deletion. This allowed us to precisely disrupt *de novo* SL synthesis in EC in a temporally precise manner. Our results revealed that lack of SPTLC1 protein in EC results in delayed vascular development. This phenotype was caused by reduced EC proliferation and tip cell formation in early angiogenic phases. In contrast, organotypic specialization of the developing vasculature was unaffected. This phenotype resembles loss of VEGFR2 in the endothelium (Zarkada *et al*, 2015). In addition, in the OIR model, where pathological neovascularization is driven primarily by VEGF, inactivation of SPT activity in EC resulted in larger vaso-obliterated zones and reduced vascular tuft formation, again suggesting impaired VEGF signaling. In agreement with this conclusion, a recent publication which described that the EC deletion of *Sptlc2* gene impaired VEGFR2-Akt signaling *in vitro* in adult arterial vessels (Cantalupo *et al*., 2020).

We also found that *Sptlc1* KO EC exhibited impaired cell proliferation, which is similar to the retinal EC in *Nos3* KO mice (Ha *et al*, 2016). However, EC nitric oxide synthase (eNOS) activity (encoded by *Nos3* gene) was elevated in *Sptlc2* ECKO adult mice (Cantalupo *et al*., 2020), which implies isoform-selective functions between SPTLC1 and SPTLC2. Alternatively, stage-specific function of the SPT enzyme or tissue-specific roles could explain these divergent phenotypes. Since VEGFR2 and eNOS functions are regulated by SL-rich membrane domains (such as lipid rafts or caveolae), we suggest that *de novo* synthesis of EC SL regulates retinal angiogenesis by alterations in membrane lipid domains that impact VEGF responsiveness of EC.

The decreased vascular development phenotype observed in *Sptlc1* ECKO mice does not resemble a hypersprouting phenotype nor is there down-regulation of BRB specific genes in mice that lack S1P receptors in EC (Jung *et al*., 2012) (Yanagida *et al*, 2020). Moreover, EC-selective deficiency in sphingosine kinase 1(encoded by *Sphk1* gene) did not display alterations in retinal vascular development (Nitzsche *et al*, 2021). Hence, we reason that membrane functions of SL metabolites, rather than GPCR-dependent signaling functions of S1P are involved.

Extracellular SL carried by lipoproteins and albumin can be taken up by cells and processed in lysosomes to generate sphingosine (Nilsson & Duan, 2006). SL can also be degraded extracellularly, followed by cellular uptake of sphingosine (Kono *et al*, 2006). However, it is less studied whether SL derived from the *de* novo pathway are transferred between various cells and tissues. Previous report documented that liver specific KO of *Sptlc2* reduces plasma SM and Cer while Sph, S1P and dhS1P are unaffected (Li *et al*, 2009). Our work demonstrates a distinctive pathway where EC supplies SL metabolites to the circulatory system and peripheral tissues. Loss of SPT activity in EC caused reduction in dhSph, Cer species and S1P in plasma and RBC. Moreover, SL reduction occurred rapidly and concurrently in RBC and plasma upon *Sptlc1* deletion in EC. The reduction of S1P levels in lipoprotein-free fraction of plasma but not in the HDL fraction of *Sptlc1* ECKO mice suggest that EC-derived S1P in the plasma interacts first with albumin, followed by stable interaction with ApoM on HDL. In the case of RBC, the rapid reduction of S1P levels occurs faster than half-life for RBC lifespan (∼22 days), indicating a constant and active SL supplementation from EC to RBC.

A pivotal question raised from our results is how various SL species produced in the EC are transferred into plasma and circulatory cells. It has been reported that EC contribute to plasma S1P levels via SPNS2, an S1P transporter (Hisano *et al*., 2012), which raises the possibility that reduced circulatory SL is due to alterations in EC-derived S1P. We believe that this is not the case. This is because loss of *Spns2* caused minor reduction of S1P levels in plasma but not in whole blood (Hisano *et al*., 2012) (Mendoza *et al*., 2012). In addition, RBC S1P transporter (encoded by *Mfsd2b* gene) knockout mice show S1P accumulation in RBC while plasma S1P is reduced (Vu *et al*, 2017). Our results from *Sptlc1* ECKO mice shows clear differences from those studies which characterized EC and RBC S1P transporter knockout mice. Having ruled out EC secretion of S1P as a possible mechanism, we next considered direct release of dhSph and Cer from EC into circulation, since such SL metabolites were reduced in *Sptlc1* ECKO plasma and RBC. While the existence of transporters for dhSph and Cer species have not been described, we do not believe that direct export of sphingoid bases and Cer species from EC into circulation are unlikely. However, alternative ways of transfer may exist. One possibility is through direct contact of EC with lipoproteins and/or RBC. Given that acid sphingomyelinase and ceramidase activities have been reported in the plasma and RBC (Xu *et al*, 2010)(Lopez *et al*, 2012), complex SL containing both reduced and unreduced sphingoid bases on the outer leaflet of the EC plasma membrane could be metabolized into Cer and sphingoid bases. SL species such as Sph can be taken up and further metabolized by circulating hematopoietic cells including RBC (Nguyen *et al*, 2021). An alternative mechanism is that EC release of circulating extracellular vesicles (EV) or exosomes from plasma membranes into circulation. It has been reported that Cer, SM, and glycosphingolipids are enriched in EV (Llorente *et al*, 2013)(Verderio *et al*, 2018) (Akawi *et al*, 2021). It is thus possible that EV derived from EC can bind to RBC or lipoproteins and SL can be taken up and metabolized.

In addition, we observed SL reductions in non-EC and peripheral tissues from *Sptlc1* ECKO mice. Specific organs and tissues may have different routes for SL transfer based on the unique barrier properties of the organ-specific endothelium. In the liver, EC form fenestrated capillaries with intercellular gaps and discontinuous basement membrane. This allows plasma and macroparticles, to cross EC barrier efficiently and access hepatocytes (Hennigs *et al*, 2021). SL exchange could therefore occur in the liver due to the porous nature of the liver sinusoidal endothelium. We postulate that the SL metabolic flux from EC to the plasma followed by hepatocyte uptake could explain the reduced liver SL content observed in *Sptlc1* ECKO mice. In the case of lung, pulmonary EC are non-fenestrated with tight junctions in between and serve as a semipermeable barrier separating circulation from lung interstitial spaces. To transport materials through lung capillaries, transcytosis acts as the primary route (Jones & Minshall, 2020). It is known to be mediated by caveolae, a critical vesicular structure, which are plasma membrane invaginations enriched in SL and sterols (Harder & Simons, 1997). We speculate that the SL flux in the lung could be mediated by this process which is affected by loss of EC *Sptlc1* gene. This mechanism could account for the reduced lung SL species in *Sptlc1* ECKO mice. In contrast, caveolae-mediated transcytosis is minimal in the CNS EC which effectively compartmentalizes metabolism of lipids between the circulation and the CNS parenchyma. Our results show that retina SL species were unaffected by lack of *de novo* SL biosynthesis in EC. Thus, the strict barrier function of the CNS EC ensures compartmentalization of SL metabolism between the neural retina and circulatory/vascular elements.

Finally, we show that *Sptlc*1 ECKO liver is protected against acetaminophen-induced hepatotoxicity. Increased GSH levels in *Sptlc1* ECKO liver suggests that glutathione synthesis, a primary detoxification agent of APAP, is involved in mediating the protective phenotype. Since serine is a source of glycine and cysteine used to synthesize glutathione, we speculate that lack of SPT activity in EC may allow the unused serine to be converted to GSH (Zhou *et al*, 2017). Moreover, our results show that that SL provided by EC is needed for APAP-induced hepatotoxicity. Recently, Cer species increases has been reported to be involved in drug induced liver injury models in mice (Li *et al*, 2020). Our liver lipidomic results further suggest a delayed accumulation of Cer in *Sptlc1* ECKO mice upon APAP treatment (Fig. 8 E and H), which correlates with its protective phenotype at early time points. It is worth noting that administration of Fumonisin B1 (FB1), a Cer synthase inhibitor, does not alleviate APAP-induced liver damage (Park *et al*., 2013). FB1 pretreatment reduces Cer content in the liver while accumulating dhSph, which is in contrast of what we observed in *sptlc1* ECKO liver. Future experiments with myriocin, a direct inhibitor of SPT complex, would be warranted to determine the role of dhSph during APAP-induced liver injury.

In conclusion, our studies suggest that *de novo* SL synthesis pathway in EC is needed for normal and pathological angiogenesis. Secondly, they also reveal a novel function of the endothelium as a source of SL metabolites in circulation and non-CNS tissues. Such mechanisms may be involved in EC dysfunction, a common mechanism in many cardiovascular and cerebrovascular diseases.

## Material and Methods

### Mouse Strains

*Sptlc1*^fl/fl^ mice were generated as previously described (Alexaki *et al*, 2017). Endothelial specific deletion of *Sptlc1*(*Sptlc1* ECKO) were generated by intercrossing *Sptlc1*^fl/fl^ mice with *Cdh5*-Cre-ERT2 strain (Sorensen et al., 2009). *Sptlc1* ECKO mice used in the experiments were *Sptlc1*^fl/fl^ carrying one *Cdh5*-ERT2-Cre allele. Controls were littermate *Sptlc1*^fl/fl^ mice. To identify *Sptlc1*^fl/fl^ allele and WT, the following primers were used: 5’-GGG TTC TAT GGC ACA TTT GGT AAG-3’(forward primer), and 5’-CTG TTA CTT CTT GCC AGT GGA C-3’ (reverse primer), generating products of 350 bp for WT and 425 bp for *Sptlc1*^fl/fl^ allele (Alexaki *et al*., 2017). The Cdh5-Cre-ERT2 allele was detected by PCR with the following primers: 5’-X-3’ (forward primer), and 5’-X-3’ (reverse primer), generating a product of 550 bp. The recombined *Sptlc1* knock-out allele was identified by PCR using 5’-CAG AGC TAA TGG AAA GGT GTC-3’ (forward primer) and 5’-CTG TTA CTT CTT GCC AGT GGA C-3’ (reverse primer), generating a product of 315 bp. To induce the deletion of *Sptlc1* in Figure 1, 3-5, 6A-C, 7 and 8, 4- to 6-week-old mice were given 2 mg tamoxifen (Sigma-Aldrich) dissolved in corn oil (Sigma) via intraperitoneal injection for 3 consecutive days. For Figure 2 to 4 and 6 D-J, neonates from *Sptlc1*ECKO strain were given 50 μg tamoxifen via intraperitoneal injection for 3 consecutive days from P1 to P3. Both male and female neonates were used for the experiments. All animal experiments were approved by the Boston Children’s Hospital Institutional Animal Care and Use Committee (IACUC).

### Mouse EC Isolation

Mouse lung EC were isolated from three to four pairs of lungs dissected from control or *Sptlc1* ECKO mice 4 weeks post tamoxifen treatment. Freshly isolated lung tissues were minced with scissors and allowed to digest at 37 °C in Liberase (0.6 U/mL, Sigma) in HBSS for 45 mins. Samples were further subjected to mechanical disruption by gentleMACS™ Octo Dissociator (Miltenyi Biotec) and filtration through fine steel mesh (40um, Falcon). Cells were washed twice with HBSS containing 0.5% Fatty Acid Free BSA. RBC were removed by ACK lysis buffer. Cells were incubated with CD45 micro beads (Miltenyi Biotec) at 4°C for 15 mins to deplete CD45+ cells. Remaining cells were incubated with CD31 micro beads (Miltenyi Biotec) at 4°C for 15 mins to enrich EC. The purity of each population was determined by flow cytometry.

Cells were stained with stained with phycoerythrin (PE)-conjugated anti-mouse CD31 (Biolegend, San Diego, CA), allophycocyanin (APC)-conjugated anti-mouse CD45 (Biolegend) in blocking solution (0.25% FAF-BSA in HBSS) with anti-CD16/32 (2.5 µg/mL) for 15 minutes on ice. Stained cells were analyzed by BD FACSCalibur. CD31 and CD45 were gated as shown in Fig. 1B.

### Western Blotting

Isolated cells from EC isolation procedure were extracted with lysis buffer (50 mM, Tris·HCl (pH 7.4), 0.1 mM EDTA, 0.1 mM EGTA, 1% Nonidet P-40, 0.1% sodium deoxycholate, 0.1% SDS, 100 mM NaCl, 10 mM NaF, 1 mM sodium pyrophosphate, 1 mM sodium orthovanadate, 25 mM sodium β-glycerophosphate, 1 mM Pefabloc SC, and 2 mg/mL protease inhibitor mixture (Roche Diagnostics)). Protein concentration was determined by Dc Protein Assay kit (Bio-Rad). 20-50µg of proteins were analyzed by SDS/PAGE and transferred onto a nitrocellulose membrane. Blots were blocked with 5% non-fat milk in TBSt. Blocked blots were incubated with primary antibodies including SPTLC1 (Santa Cruz; 1:1000), eNOS (BD Biosystem; 1:2000) β-actin (Sigma; 1:5000) at 4°C overnight. Blots were washed 3 times with TBSt for 5 mines before adding the secondary antibodies. After another 3 times wash with TBSt, blots were developed with ECL western blotting substrate (Pierce) and imaged by Azure 600 imaging system (Azure Biosystems).

### Sphingolipid Measurements of Isolated Cells

Isolated cells from EC isolation procedure were counted and 2-5×10^6^ cells were subjected to HPLC-mass spectrometry (MS/MS) by the Lipidomics Core at the Stony Brook University Lipidomic Core (Stony Brook, NY) using TSQ 7000 triple quadrupole mass spectrometer, operating in a multiple reaction monitoring-positive ionization mode as described (Bielawski *et al*, 2010).

### Retina Immunohistochemistry in Flat Mounts

Eyeballs from Ctrl and *Sptlc1* ECKO mice were enucleated and fixed in 4% PFA in PBS at room temperature for 20 min. Retinas were isolated under dissection microscopy, permeabilized in 0.5% Triton X-100 in PBS at room temperature for 30 min and blocked with 1% Bovine Serum Albumin (BSA) in PBSt at room temperature for 30 mins. Primary antibodies were added to the sections with indicated dilution in PBSt (NG2 (Sigma, 1:200), Phsopho-histone H3 (BioLegend, 1:400), ESM1 (R and D system, 1:200), LEF1 (Cell Signaling Technology, 1:200), TFRC (Novus Biologicals, 1:200), CLDN5 (Invitrogen, 1:200), MFSD2A (A kind gift from David Silver, 1:200), Alexa Fluor 647 isolectin GS-IB4 conjugate (Invitrogen, 1:500), Alexa Fluor 488-conjugated rabbit monoclonal anti-ERG (Abcam, 1:200), Alexa Fluor 647-conjugated rabbit monoclonal anti- ERG (Abcam, 1:200), Cy3- conjugated mouse monoclonal anti-a-smooth muscle actin (Sigma, 1:500)) at 4°C overnight. Retinas were further washed 3 times with PBS for 10 mins at room temperature and incubated with fluorescent conjugated secondary antibodies with 1:500 dilution in PBSt for 2 hours at room temperature. Stained retinas were washed 3 times with PBS for 10 mins at room temperature and mounted using Fluoromount-G slide mounting medium (Southern Biotech). Flat Mount Retina Slides were proceeded to image with confocal microscopy (Zeiss).

### Oxygen Induced Retinopathy

Retinal neovascularization was observed using the OIR ((Smith *et al*., 1994)). In brief, neonates together with their mother were exposed to 75% oxygen from P7 to P12. Upon returning to room air (20% O2) at P12 the relative hypoxia induced neovascularization over the period P12 through P17 in these mice, with P17 as the maximum of neovascularization. For experiments in Figure 4H-K, 100μg tamoxifen were administrated via intraperitoneal injection at P12. Eyeballs were collected at P10, P12 or P17 for immunofluorescence (See section above). Retinal neovascularization and vaso-obliteration were quantified using Image J as previously reported (Stahl *et al*, 2009).

### Sphingolipid Measurements of Liver and Lung

Tissues were homogenized in 20mM Tris-HCl pH 7.5 with tissue homogenizer (Fischer Scientific). Protein concentration was determined by Dc Protein Assay kit (Bio-Rad). 2-6 mg of total protein were used for HPLC-MS/MS by the Lipidomics Core at the Stony Brook University Lipidomic Core (https://osa.stonybrookmedicine.edu/research-core-facilities/bms Stony Brook, NY) using TSQ 7000 triple quadrupole mass spectrometer, operating in a multiple reaction monitoring-positive ionization mode as described (Bielawski *et al*., 2010)..

### Plasma S1P Measurement

Plasma S1P and RBC S1P in Figure 6A-C was extracted as previously described (Frej *et al*, 2015) with minor modification. Plasma aliquots (5 or 10 mL) were first diluted to 100 mL with TBS Buffer (50 mM Tris-HCl pH 7.5, 0.15 M NaCl). S1P was extracted by adding 100 mL precipitation solution (20 nM D7-S1P in methanol) followed by 30 s of vortexing. Precipitated samples were centrifuged at 18,000 rpm for 5 min and supernatant were transferred to vials for UHPLC-MS/MS analysis (see below). C18-S1P (Avanti Lipids) was dissolved in methanol to obtain a 1 mM stock solution. Standard samples were prepared by diluting the stock in 4% fatty acid free BSA (Sigma-Aldrich) in TBS to obtain 1 mM and stored at −80°C. Before analysis, the 1 mM S1P solution was diluted with 4% BSA in TBS to obtain the following concentrations: 0.5 mM, 0.25 mM, 0.125 mM, 0.0625 mM, 0.03125 mM, 0.0156 mM, and 0.0078 mM. S1P in diluted samples (100 mL) were extracted with 100 mL of methanol containing 20 nM of D7-S1P followed by 30 s of vortexing. Precipitated samples were centrifuged at 18,000 rpm for 5 min and the supernatants were transferred to vials for mass spectrometric analysis. The internal deuterium-labeled standard (D7-S1P, Avanti Lipids) was dissolved in methanol to obtain a 200 nM stock solution and stored at −20°C. Before analysis, the stock solution was diluted to 20 nM for sample precipitation. The samples were analyzed with Q Exactive mass spectrometer coupled to a Vanquish UHPLC System (Thermo Fisher Scientific). Analytes were separated using a reverse phase column maintained at 60°C (XSelect CSH C18 XP column 2.5 mm, 2.1 mm X 50 mm, Waters). The gradient solvents were as follows: Solvent A (water/methanol/formic acid 97/2/1 (v/v/v)) and Solvent B (methanol/acetone/water/formic acid 68/29/2/1 (v/v/v/v)). The analytical gradient was run at 0.4 mL/min from 50–100% Solvent B for 5.4 min, 100% for 5.5 min, followed by one minute of 50% Solvent B. A targeted MS2 strategy (also known as parallel reaction monitoring, PRM) was performed to isolate S1P (380.26 m/z) and D7-S1P (387.30 m/z) using a 1.6 m/z window, and the HCD-activated (stepped CE 25, 30,50%) MS2 ions were scanned in the Orbitrap at 70 K. The area under the curve (AUC) of MS2 ions (S1P, 264.2686 m/z; D7-S1P, 271.3125 m/z) was calculated using Skyline (MacLean et al., 2010). Quantitative linearity was determined by plotting the AUC of the standard samples (C18-S1P) normalized by the AUC of internal standard (D7-S1P); (y) versus the spiked concentration of S1P (x). Correlation coefficient (R2) was calculated as the value of the joint variation between x and y. Linear regression equation was used to determined analyte concentrations.

### Plasma and RBC Collection

For non-terminal blood collection, blood was collected from submandibular vein via cheek punch. For terminal blood collection, mice were euthanized with CO_2_. Blood was recovered via vina cava. Blood samples were collected in tubes containing 1-5 ul 0.5 M EDTA depending on sample volume. Samples were centrifuged at 2,000g for 10 min. Plasma samples were collected and stored at −80°C. RBC were washed twice with HBSS and cell number was counted by hemocytometer. 1010 cells were pelleted by centrifugation at 2,000g for 10 mins and store at −80°C.

### Sphingolipid Measurement in Plasma and Retina

Retinas were placed on dry ice for their extraction. Afterwards, 600 µl of ice cold (Freezer for 1 hour, kept on ice while processing outside) LC-MS methanol was added to each sample, followed by 10 µl of the deuterated sphingolipid labeled internal standards. Samples were then vortexed and allowed to equilibrate on ice for 15 minutes. This was followed by two 15 minutes sonication steps in an ice bath. Next, samples were centrifuged at 12000 *g* for 15 minutes at 6⁰C. Finally, 120 µl of the supernatant were transferred to an LC-MS amber vial equipped with a 150 µl insert for LC-MS injection. A pool of 25 µL of all samples was used as QC of the injection. Samples were injected on an Acquity UPLC system coupled to a Xevo TQ-S mass spectrometer (Waters, Milford, MA) as previously described (Akawi *et al*., 2021).For plasma analyses, samples were thawed at 4⁰C in the refrigerator. Then, samples were extracted as previously described (Akawi *et al*., 2021). Briefly, a volume of 10 µl of the sphingolipid internal standard mix containing labeled internal standards. Samples were then vortexed for 10 seconds and equilibrated with the internal standard at room temperature. Then, a volume of 250 µl LC-MS methanol was added to each sample, followed by a 10 seconds vortex period and sonication for 15 minutes in an ultrasound bath with ice. Next, samples were centrifuged at 12000 *g* for 15 minutes at 6⁰C. Finally, 120 µl of the supernatant were transferred to an LC-MS amber vial equipped with a 150 µl glass insert. A pool of 25 µL of all samples was used as QC of injection. Samples were then analyzed using the same method as described for the retinas.

### APAP-induced Liver Injury

Mice were fasted for 16hrs. APAP was dissolved in PBS and administered to the mice via intraperitoneal injection (300mg/kg). Food and water were given ad lib after treatment. Plasma and liver samples were collected at indicated time for further analysis. Liver function was measured using AST activity assay kits (Sigma) according to the manufacturer’s instructions. Liver GSH levels were measured using glutathione assay kits (Cayman Chemical) according to the manufacturer’s instructions.

### Confocal Microscopy and Image Processing

Images were acquired using a LSM810 confocal microscope (Zeiss) equipped with an EC Plan-Neofluar 10x/0.3, a Plan-Apochromat 20x/0.8 or a Plan-Apochromat 40x/1.4 Oil DIC objective. Images were taken using Zen2.1 software (Zeiss) and processed and quantified with Fiji (NIH). Figures were assembled using Adobe Affinity Photo and Windows Office Powerpoint.

### Statistics

For datasets containing exactly two groups, an unpaired two-tailed Student’s t test was used to determine significant differences. For datasets containing exactly four groups, one-way ANOVA followed by Bonferroni’s post hoc test was used to determine significant differences. P value less than 0.05 was statistically significant.

## Acknowledgements

This work was supported in part by the NIH grants (R35-HL135821 and R01EY031715 to TH), Intramural Research Programs of the National Institutes of Health, National Institute of Diabetes and Digestive and Kidney Disease to RLP, National Institutes of Health, and American Heart Association Postdoctoral Fellowship (18POST33990339) to AK.

## Supplemental Figure Legends

**Figure 1 Supplement 1.**
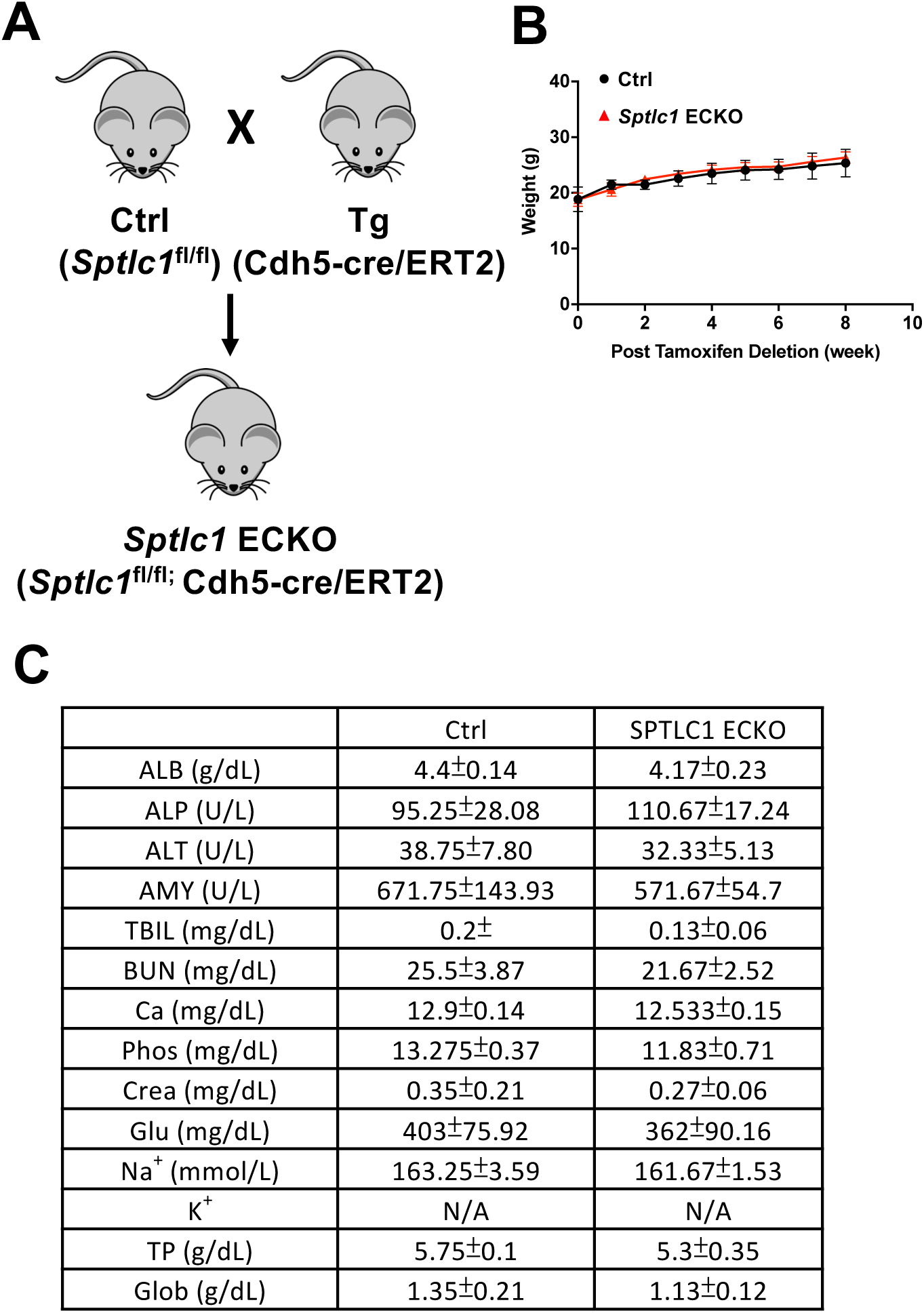
*Sptlc1* ECKO mice exhibit similar body weight and blood chemistry as control mice. (A) schematic diagram illustrates generation of *Sptlc1* ECKO mice. (B) Body weight was measured at indicated time after tamoxifen injection in Ctrl and *Sptlc1* ECKO mice. (C) Blood chemistry analysis was performed in Ctrl and *Sptlc1* ECKO serum. Data are expressed as mean±SD.

**Figure 1 Supplement 2.**
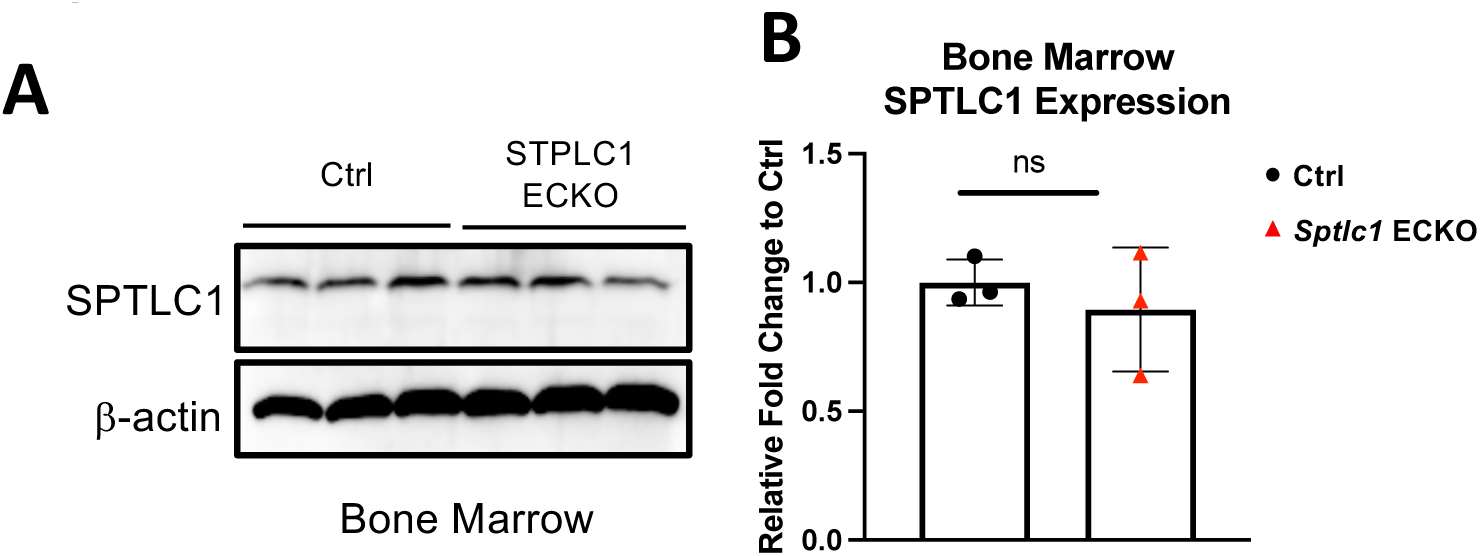
SPTLC1 Expressions are not affected in bone marrow of *Sptlc1* ECKO mice. (A) Bone marrow cells isolated from Ctrl and *Sptlc1* ECKO mice were analyzed by Western Blotting for SPTLC1 expressions. (B) Relative fold change to Ctrl samples were quantified by ImageJ software. Data are expressed as mean±SD. ns, nonsignificant.

**Figure 1 Supplement 3.**
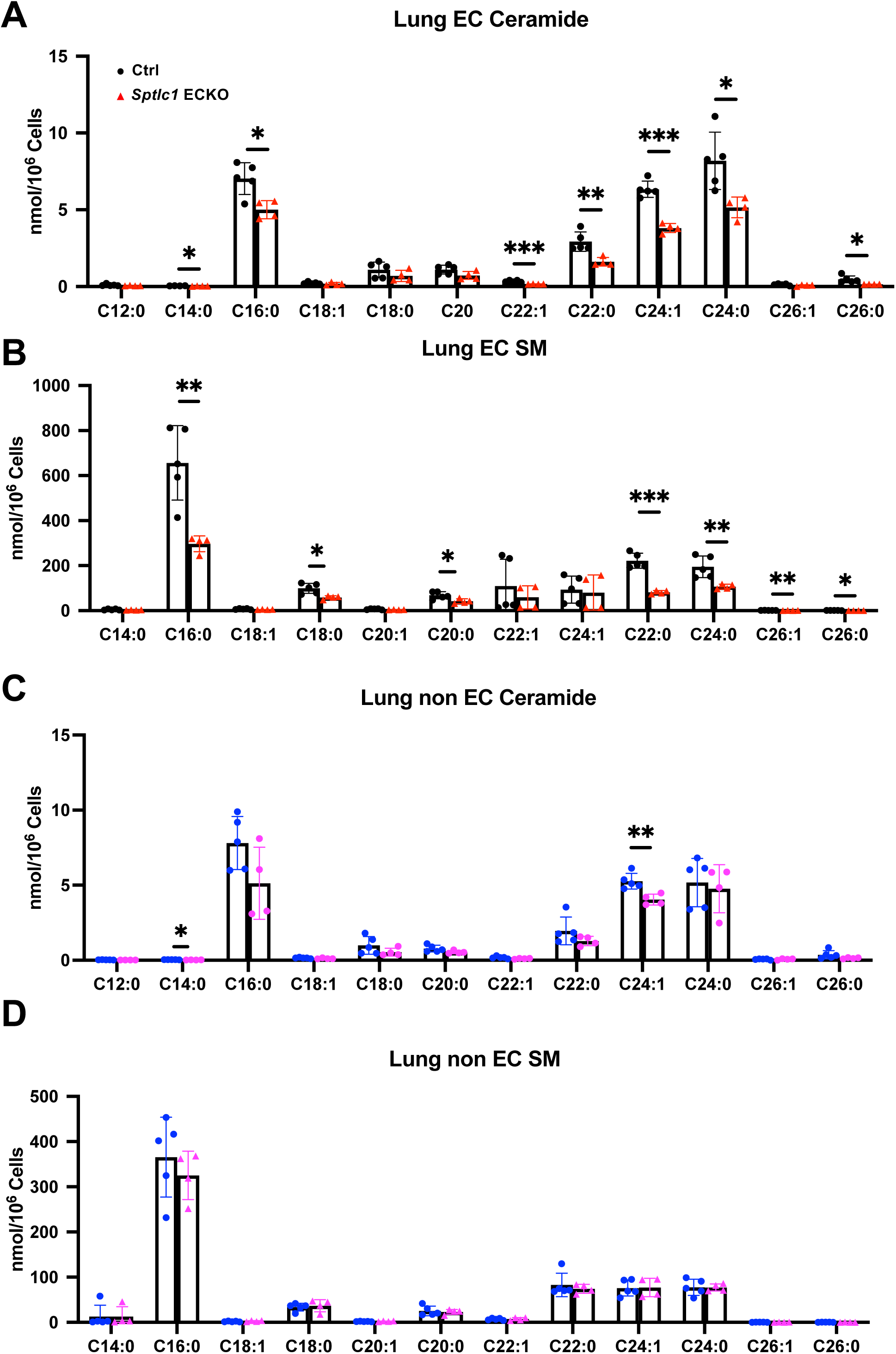
Cer and SM are reduced in *Sptlc1* KO EC while they are mildly affected in non-EC. (A) Lung EC Cer with different chain lengths in Ctrl and *Sptlc1* ECKO mice were determined by LC-MS/MS. (B) Lung EC SMs with different chain length in Ctrl and *Sptlc1* ECKO mice were determined by LC-MS/MS. (C) Lung non-EC Cer with different chain length in Ctrl and *Sptlc1* ECKO mice were determined by LC-MS/MS. (B) Lung non-EC SMs with different chain length in Ctrl and *Sptlc1* ECKO mice were determined by LC-MS/MS. Data are expressed as mean±SD. *p<0.05. **p<0.01. ***P<0.001. ns, nonsignificant.

**Figure 2 Supplement 1.**
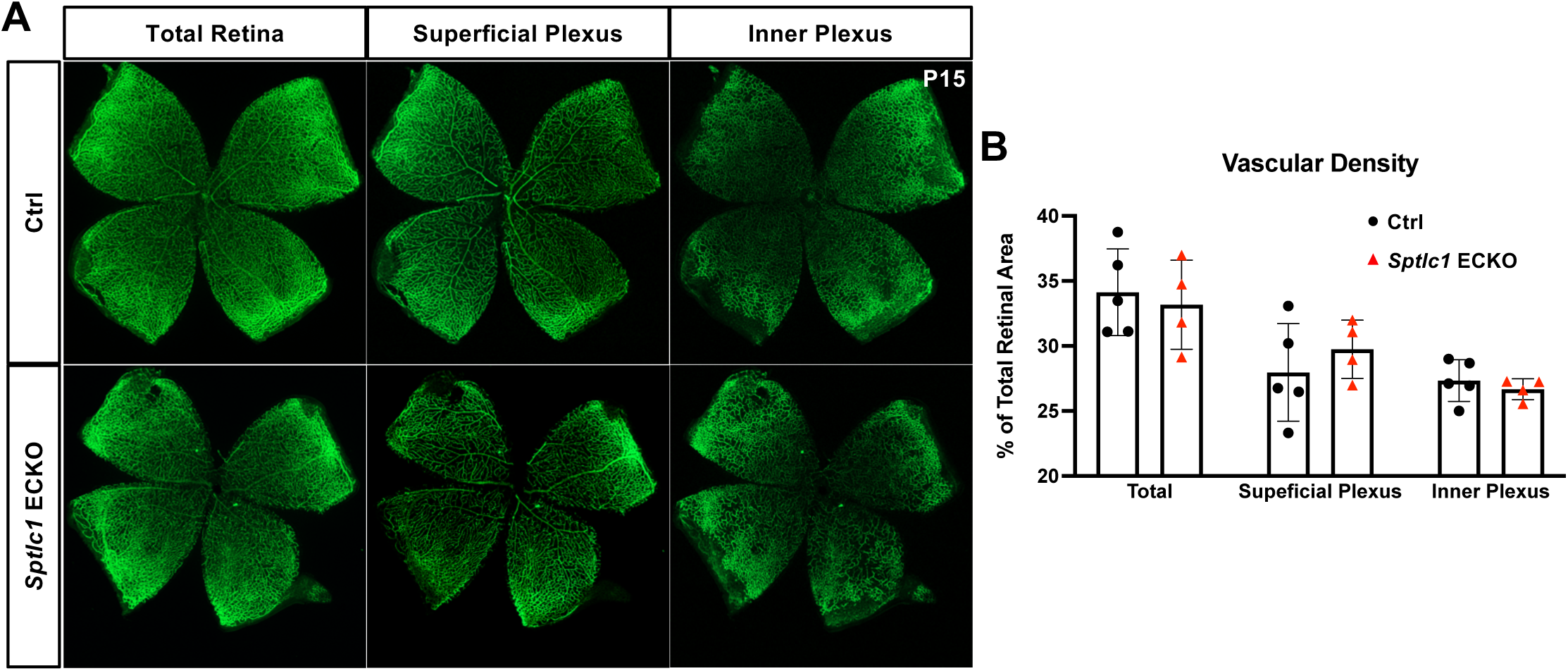
Retinal vascular development is normalized at P15 in *Sptlc1* ECKO mice. (A) Retinal vascular plexus at P15 of Ctrl and *Sptlc1* ECKO mice were immunostained with IB4. Vascular density of total vasculature, superficial plexus and inner plexus were quantified in (B). Data are expressed as mean±SD.

**Figure 6 Supplement 1.**
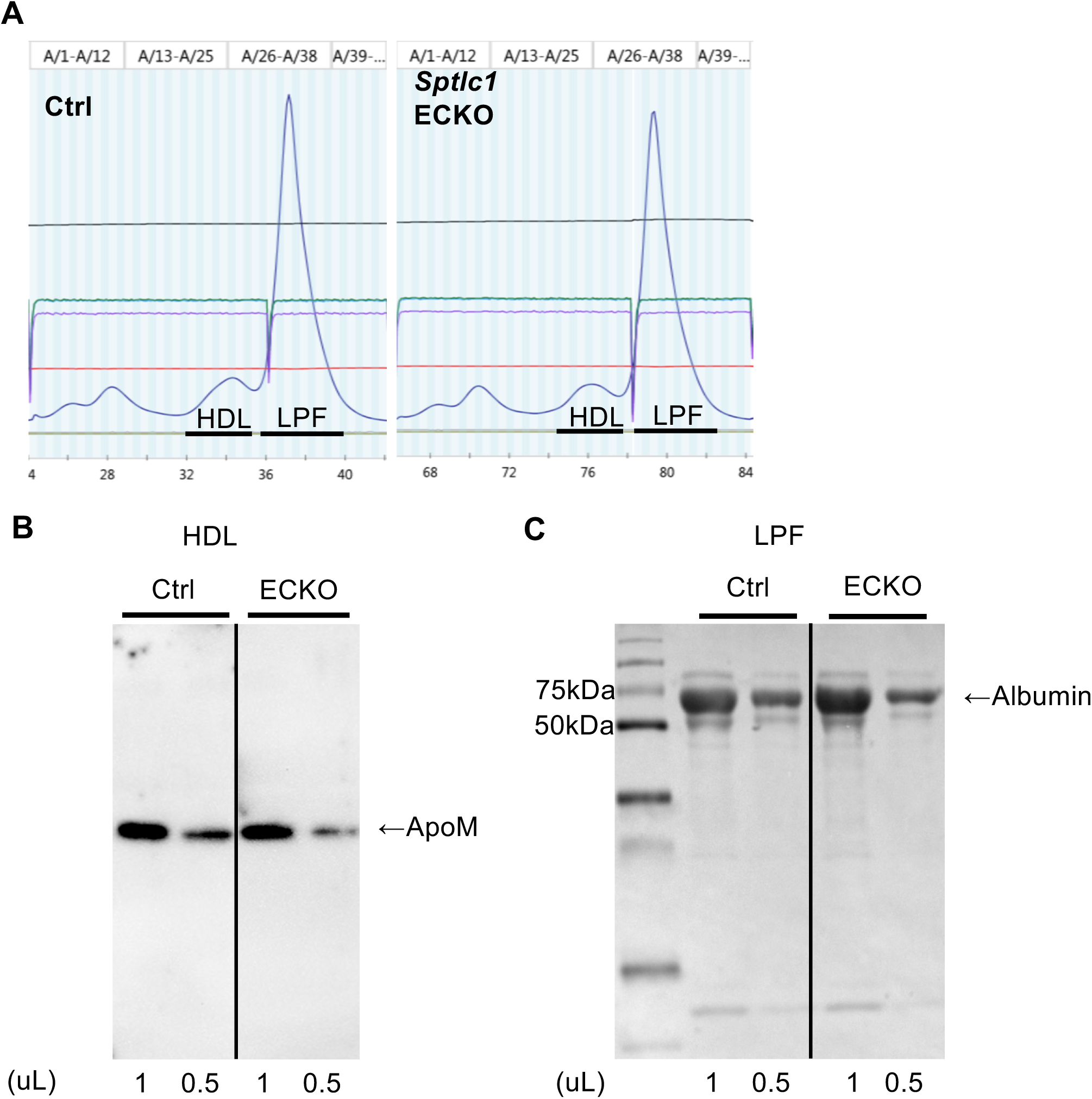
Plasma FPLC fractionation indicates similar protein abundance of ApoM and Albumin in HDL and lipoprotein free fraction between Ctrl and *Sptlc1* ECKO mice. (A) HPLC analyses of plasma samples demonstrated the abundance of proteins in HDL and lipoprotein free (LPF) fractions in Ctrl and *Sptlc1* ECKO mice as indicated by O.D.280 measurements (Blue lines). (B) HDL fraction from (A) were analyzed by Western blotting for ApoM with indicated volume of sample loaded. (C) LPF fraction of plasma from (A) were analyzed by SDS-PAGE followed by Coomassie staining with indicated volume of sample loaded. Albumin is identified based on molecular weight.

**Figure 6 Supplement 2.**
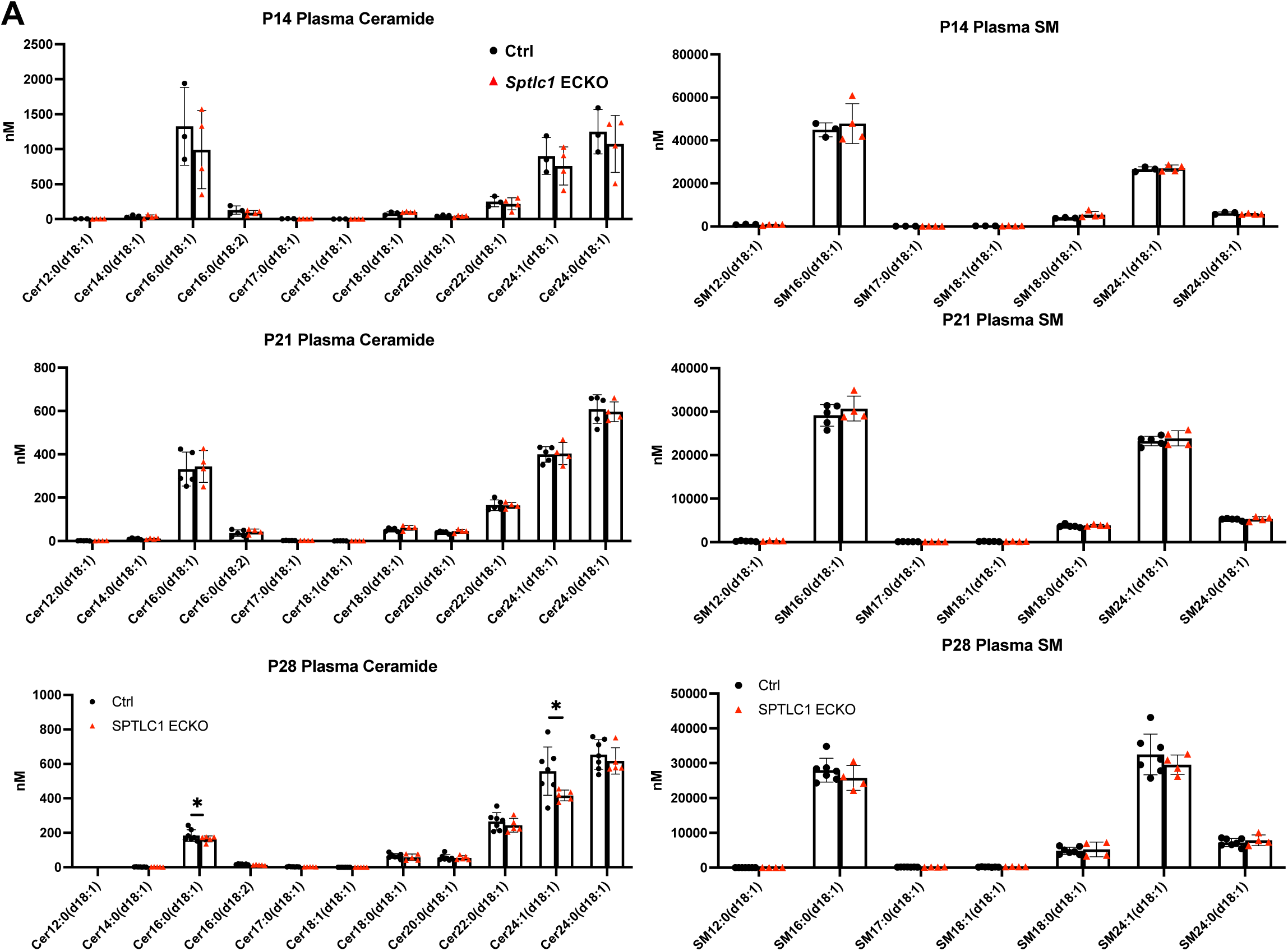
Plasma Cer and SM are not affected in postnatal deletion of *Sptlc1* ECKO mice. (A) and (B) P14 Cer and SM with different chain length in Ctrl and *Sptlc1* ECKO mice were determined by LC-MS/MS. (C) and (D) P21 Cer and SM with different chain length in Ctrl and *Sptlc1* ECKO mice were determined by LC-MS/MS. (E) and (F) P28 Cer and SM with different chain length in Ctrl and *Sptlc1* ECKO mice were determined by LC-MS/MS. Data are expressed as mean±SD. *p<0.05

